# The NSL complex mediated nucleosome landscape is required to maintain transcription fidelity and suppression of transcription noise

**DOI:** 10.1101/419408

**Authors:** Kin Chung Lam, Ho-Ryun Chung, Giuseppe Semplicio, Vivek Bhardwaj, Shantanu S. Iyer, Herbert Holz, Plamen Georgiev, Asifa Akhtar

## Abstract

Nucleosomal organization at gene promoters is critical for transcription, with a nucleosome-depleted region (NDR) at transcription start sites (TSSs) being required for transcription initiation. How NDR and the precise positioning of the +1 nucleosome is maintained on active genes remains unclear. Here, we report that the *Drosophila* Non-Specific Lethal (NSL) complex is necessary to maintain this stereotypical nucleosomal organization at promoters. Upon NSL1 depletion, nucleosomes invade the NDRs at TSSs of NSL-bound genes. NSL complex member NSL3 binds to TATA-less promoters in a sequence-dependent manner. The NSL complex interacts with the NURF chromatin remodeling complex and is necessary and sufficient to recruit NURF to target promoters. The NSL complex is not only essential for transcription but is required for accurate TSS selection for genes with multiple TSSs. Further, loss of NSL complex leads to an increase in transcriptional noise. Thus, the NSL complex establishes a canonical nucleosomal organization that enables transcription and determines TSS fidelity.

## Introduction

Chromatin structure and organisation are fundamental to the regulation of gene transcription. Chromatin at active gene promoters is characterized by a distinct nucleosomal organization (Lai and Pugh, 2017). Transcriptional start sites (TSS) are embedded in a nucleosome depleted region (NDR), which enables pre-initiation complex formation (Workman and Roeder, 1987). The NDR is bordered by the well-positioned +1 nucleosome followed by a regular array of nucleosomes. This organization is thought to be required for transcription initiation (Lai and Pugh, 2017).

The standard model of transcription initiation is based on genes that are activated in a tissue-specific manner (Kadonaga, 2012). The current model supports that TBP binds to the TATA box at promoters of tissue-specific genes, where the assembly of RNA polymerase II (Pol II) is initiated and TFIIA, B, D, E, F, and H form the preinitiation complex. TBP-TATA box binding occurs at around 30 bp upstream of the TSS and thus defines a sharp and precise transcription start site (Carninci et al., 2006; Ni et al., 2010; Yamamoto et al., 2009). However, this model does not represent the majority of genes, as multicellular organisms express a range of housekeeping genes that are critical for homeostatic maintenance. Unlike tissue-specific genes, housekeeping genes have highly dispersed TSSs that are scattered over up to 100 bp (Carninci et al., 2006; Ni et al., 2010). This difference in transcription initiation patterns between housekeeping and tissue-specific gene promoters are conserved across species including fish, flies and mammals (Carninci et al., 2006; Haberle et al., 2014; Ni et al., 2010).

The precise selection of TSS is dependent on the tissue and developmental stage (Haberle et al., 2014) and thus pose an important aspect of transcription regulation as changes in TSS can affect RNA stability and the resulting protein isoforms. Despite its importance, the nature and the causative relationship of the DNA sequence and transcription factors that direct TSS selection at housekeeping genes remains poorly understood. Compared to focused promoters of tissue-specific genes, dispersed housekeeping gene promoters contain distinct sets of core promoter motifs and binding proteins (Vo Ngoc et al., 2017). Dispersed promoters generally lack a TATA box or Inr elements, but rather contain motif1, 6, 7 and DRE (Rach et al., 2009; Vo Ngoc et al., 2017). While the TATA box initiates the binding of TBP and then Pol II, it is still not clear what instructs Pol II to initiate transcription at the dispersed promoters in the absence of a TATA box or Inr elements. Likewise, distinct set of proteins are found on dispersed promoter: Motif 1 binding protein (M1BP) recognizes motif 1 and DREF binds to DREs. Therefore, they are believed to be binding to the dispersed promoters only. The difference in DNA motifs, protein binding and transcription patterns between dispersed housekeeping and focused promoters argues for fundamental differences in the mechanisms of transcription initiation.

The distinction between the two major types of promoters could be a result of chromatin modifying factors, which influence the local chromatin modifications and organization. Elegant work in yeast, flies and mammals has demonstrated the importance of chromatin remodeling complexes in nucleosome organization (Alkhatib and Landry, 2011; Lai and Pugh, 2017; Struhl and Segal, 2013). Chromatin remodeler complexes can be broadly classified into four families (ISWI, CHD/Mi- 2, INO80/SWR1, and SWI/SNF) based on the protein domains of their catalytic ATPase subunits. Each remodeler has its characteristic molecular structure, target genomic locations and roles in cells. In higher eukaryotes, it is not yet clear which trans-acting factors are responsible for the nucleosomal organization at TSSs. How chromatin remodeling complexes work in concert with other chromatin modifying enzymes and transcription machinery to facilitate the transcription process remains an active area of research (Lai and Pugh, 2017; Struhl and Segal, 2013).

The Drosophila Non-Specific-Lethal (NSL) complex is a chromatin modifying complex. It contains the histone H4 lysine 16 acetyl transferase MOF, as well as NSL1, NSL2, NSL3, MCSR2, MBDR2, Z4, Chromator and WDS (Mendjan et al., 2006; Raja et al., 2010). Underpinning its importance, loss of NSL complex members leads to lethality during early development in flies (Raja et al., 2010), while heterozygous mutations in NSL1 and NSL2 orthologues *KANSL1* and *KANSL2* underlie intellectual disability in humans (Gilissen et al., 2014; Koolen et al., 2012; Zollino et al., 2012). The NSL complex binds to the dispersed housekeeping gene promoters and this feature is remarkably conserved from flies to human (Chelmicki et al., 2014; Lam et al., 2012; Ravens et al., 2014). However, the mechanism by which the NSL complex regulates housekeeping gene expression remains unknown. Therefore, studying how the NSL complex functions is an important paradigm to understand how transcription factors specifically target the vast number of housekeeping genes and mediate transcription in a way that is fundamentally different from what we typically associate with tissue-specific or developmental genes.

Here, we report that the NSL complex is necessary to maintain the stereotypical nucleosomal organization at promoters. Upon NSL1 depletion, nucleosomes invade the NDR at the TSS of NSL-bound genes. We also uncover that binding of the NSL complex to TATA-box-less housekeeping gene promoters is directed by AT-rich sequences. Accordingly, we can predict the in vivo NSL complex binding by AT-rich sequences and chromatin context. Mechanistically, we show that the NSL complex recruits the NURF complex to maintain the nucleosome pattern that is typical of dispersed promoters. This nucleosome pattern is important for gene regulation as its disruption leads to spurious transcription start site selection and increase in transcriptional noise. Our data illustrate how housekeeping gene promoters can be targeted by the NSL complex, which then impose a specific nucleosome pattern, TSS selection and transcription noise regulation in the Drosophila genome.

## Results

### NSL complex loss leads to reduced nucleosome patterning at target TSSs

The NSL complex is an important regulator for the majority of active promoters (Chelmicki et al., 2014; Lam et al., 2012), we sought to understand its roles in establishing the chromatin landscape at promoters. In order to study if the NSL complex is required for nucleosomal organization, we knocked down (KD) NSL1, and GST (control) in S2 embryonic cells and performed MNase-seq (Figure S1A-C). Many genes showed a decrease in the nucleosome signal at +1 nucleosome position upon NSL depletion, most pronounced at promoter regions (Figure 1B, 1C). Nucleosome occupancy changed around NSL-bound, but not around NSL-non-bound promoters (Figure 1C). We observe a decrease in the occupancy of the +1 nucleosome, concomitant with an increased occupancy at the NDR, indicating an invasion of the +1 nucleosome into the NDR. This was supported by nucleosome profiles in control and NSL1 KD, showing an average shift of the +1 nucleosome towards the TSS (Figure 1D). Downstream of the +1 nucleosome the array also shifts towards the TSS. The mean 5’ end position of the +1 nucleosome shifts upstream towards the TSS in a NSL1 binding dependent manner (Figure 1E).

**Figure 1.**
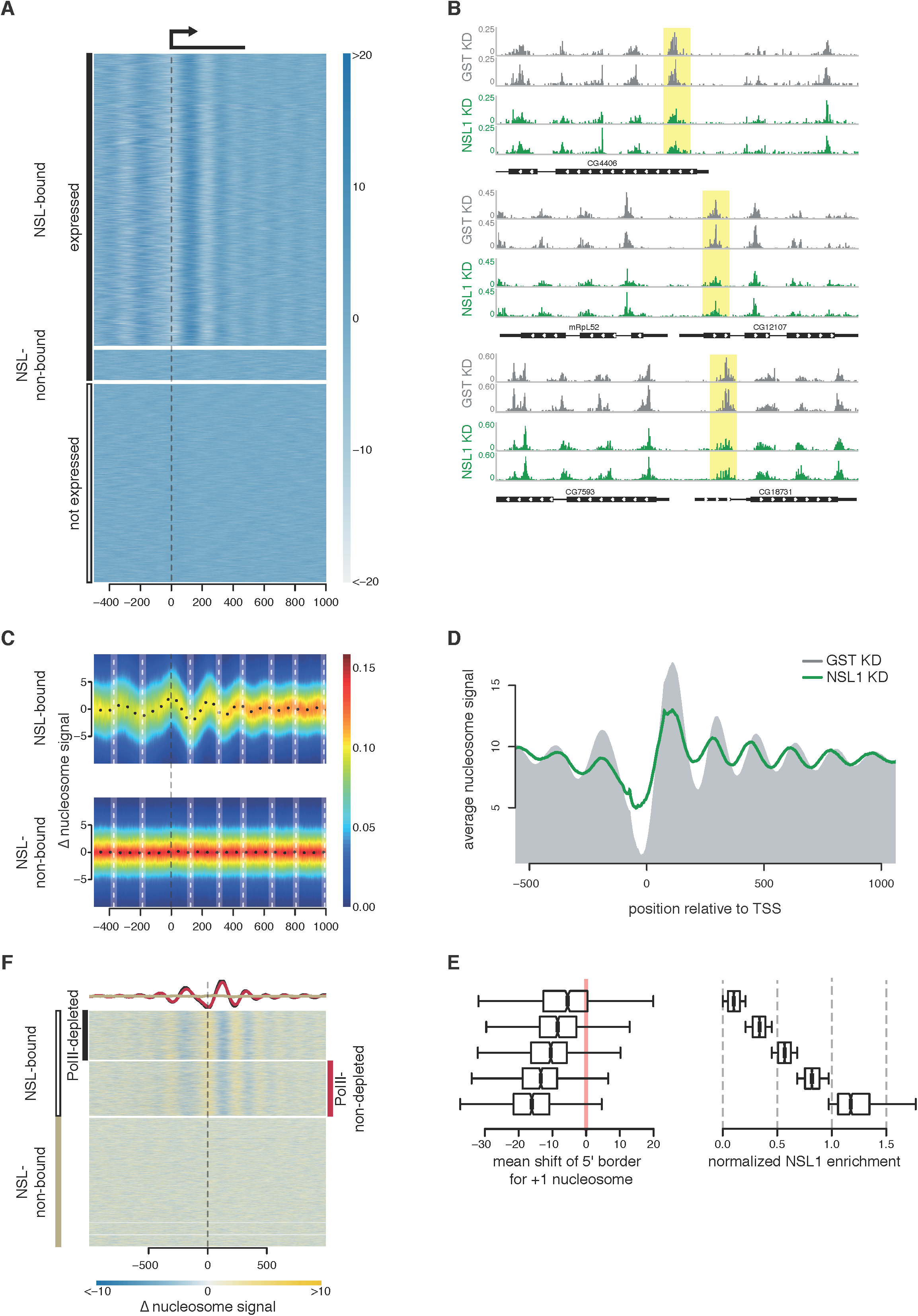
NSL complex loss leads to reduced nucleosome patterning at target TSSs. (A) Heatmap showing nucleosome signal on NSL-bound active genes (top), NSL-non-bound active genes (middle) and inactive genes (bottom), −500 bp to and +1,000 bp of the TSS. (B) Representative examples of nucleosome density (read counts normalized to sequencing depth) in control (grey) and NSL1 KD (green). The +1 nucleosomes are shaded in yellow. (C) Heatmap depicting the difference in nucleosome densities in control and NSL1 KD −500 bp to +1,000 bp of TSS of NSL-bound genes (top) and NSL-non-bound genes (bottom). The y-axis represents changes in nucleosome signal. The white vertical dashed lines denote WT nucleosome positions. The color scale bar indicates scatter density. (D) Summary plot showing the nucleosome positions in wild-type (grey shaded area) and NSL1- depleted cells (green line). (E) Quantification of the shift in the +1 nucleosomes in base pairs (right) for genes with different NSL1 log2 ChIP/Input (left). (F) Heatmap showing changes in nucleosome density upon NSL1 KD for NSL-bound genes with Pol II loss upon NSL1 KD (top), NSL-bound genes with no Pol II loss (middle) and NSL- non-bound genes (bottom). The summary plot depicts the average nucleosome density of NSL- bound genes with Pol II loss (black) and NSL-bound genes without Pol II loss (red).

The NSL complex is required for the recruitment of RNA Polymerase II (Pol II) (Lam et al., 2012), thus the changes in the nucleosomal organization could be a secondary consequence of the loss of Pol II. To address this issue, we categorized genes into three groups: (1) NSL-bound with Pol II loss; (2) NSL-bound without Pol II loss upon NSL1 KD; and (3) genes not targeted by NSL. The changes in the nucleosomal organization correlated with NSL binding (groups 1 and 2) and were present irrespective of Pol II loss. Group 3 genes failed to show significant changes (Figure 1F). Thus, the NSL complex affects the nucleosomal organization at promoters independent of the changes in Pol II recruitment.

### The NSL complex recruits the NURF complex and maintains nucleosome pattern at promoters

As NSL complex members have no chromatin remodeling activity reported to date, we asked whether they function with chromatin remodelers to position nucleosomes at promoters (Figure S2A). We used a Gal4-NSL3 reporter system (Lam et al., 2012; Raja et al., 2010), where tethering of Gal4-NSL3 to the promoter of a UAS-driven luciferase reporter results in luciferase up-regulation in S2 cells (Figure 2A). We performed RNAi against a candidate set of chromatin remodelers (NURF301, ISWI, BRM, INO80 and CHD3) (Figure S2B). KD of NURF301 caused a strong reduction in luciferase activity, comparable to the reduction observed upon MOF KD. KD of ISWI led to a milder decrease. KD of BRM caused a strong decrease in luciferase activity. However, the protein levels of MOF are severely reduced upon BRM KD, which was not true in NURF301 and ISWI KDs (Figure S2B). In contrast, INO80 and CHD3 KDs did not attenuate NSL3-mediated activation. Thus, the NURF complex is required for NSL3-mediated transcription activation.

**Figure 2.**
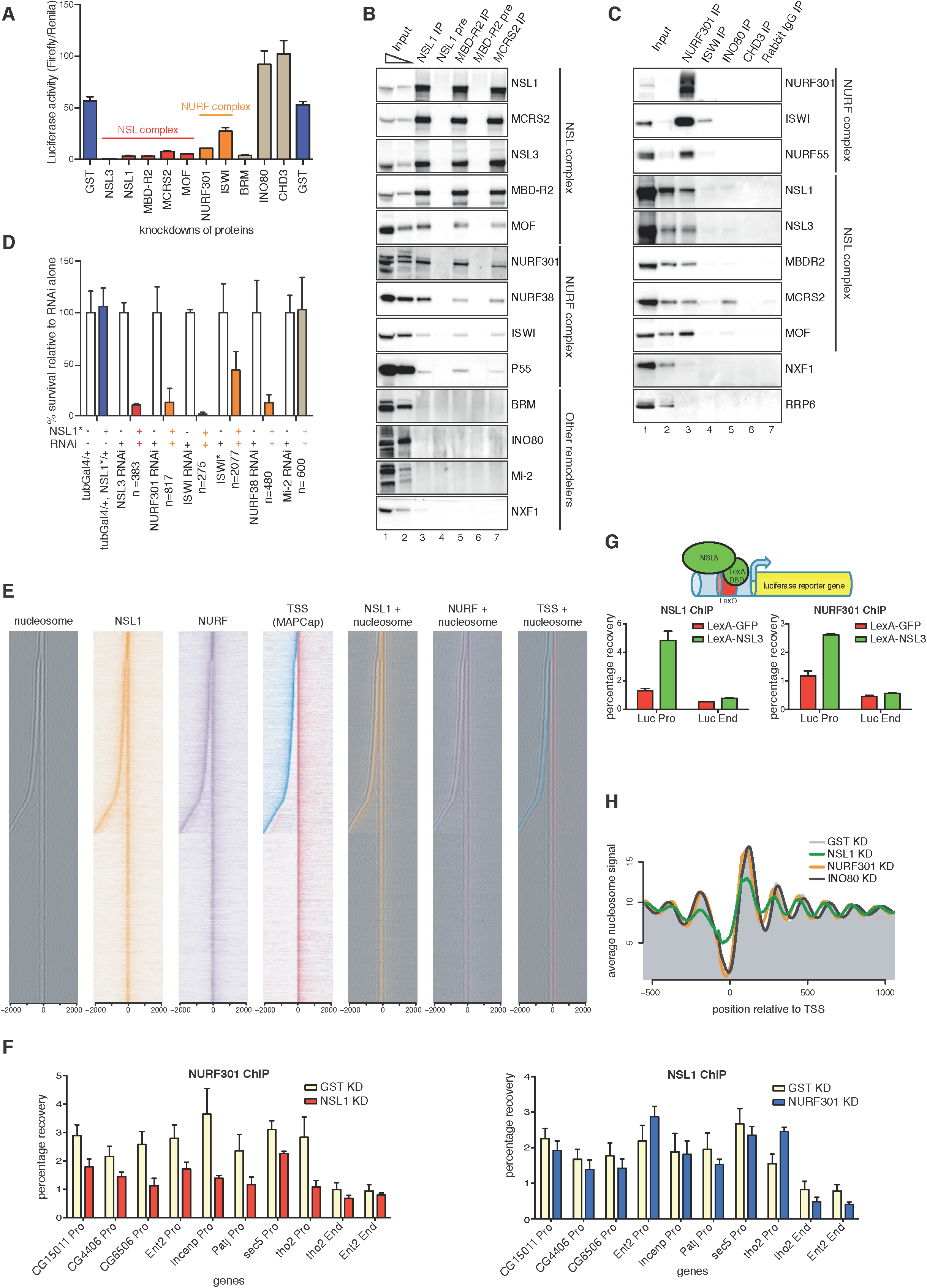
The NSL complex recruits the NURF complex and maintains nucleosome pattern at promoters. (A) Luciferase activity (ratio of Firefly luciferase/Renila lucifrease) in a Gal4-NSL3 reporter system in KDs for GST (control, left- and right-most), NSL and NURF complex members, BRM, INO80 and CHD3. Error bars represent standard deviation (SD) of three independent experiments. (B) Immunoprecipitation of the endogenous NSL complex members NSL1, MBDR2 and MCRS2. Pre-immune Sera (NSL1, MBD-R2) served as controls. The Western blot was probed by antibodies against the NSL- and NURF- complex, other chromatin remodelers, as well as NXF1, a protein involved in mRNA export (as a negative control). (C) Immunoprecipitation of the chromatin remodelers NURF301, ISWI, INO80 and CHD3. Rabbit IgG served as a control. The Western blot was probed by antibodies against the NSL- and NURF-complex. INO80 immunoprecipitates MCRS2, but not other NSL complex members. NXF1 and RRP6 served as negative controls. (D) Bar chart showing of viability of flies when combining KD of NSL3, NURF301, ISWI (dominant negative mutant, marked with *, and RNAi), NURF38 or Mi-2 with heterozygous NSL1 mutant (blue bars) relative to RNAi/mutant alone (white bars). Heterozygous NSL1 mutant alone (blue bar) does not cause lethality. The mean ± s.d. of at least three independent crosses is shown. The numbers of flies counted were indicated by n. (E) Heatmaps displaying the input-normalized ChIP enrichments of NSL1 (orange) and NURF301 (purple) ± 2kb around the MAPCap TSS. Nucleosome signal is depicted in black. MAPCap TSS positions are indicated in red (Watson strand) and blue (Crick strand). Nucleosome signals are overlaid with the ChIP-seq and TSS signal (right). (F) ChIP–qPCR analyses for NURF301 (top) and NSL1 (bottom) for KDs against GST (pale yellow), NSL1 (red) and NURF301 (blue). qPCR analysis was carried out with primer sets positioned at the promoters (Pro) and ends (End) of indicated genes. Results are expressed as mean (± s.d.) of relative percentage recovery of IP material over input material. (G) Schematic: LexA-NSL3 is used to activate expression of lexO-luciferase reporter in transgenic flies. Bar charts: ChIP–qPCR experiments performed with NSL1 (left) and NURF301 (right) antibodies and primer pairs specific for promoter and end regions of the reporter gene. Results are expressed as mean (± s.d.) of relative percentage recovery of IP material over input material. (H) Summary plot depicting the nucleosome signal −500 bp to 1,000 bp of the TSS, in wild type (grey shaded area), NSL1-depleted (green), NURF301-depleted (orange) and INO80-depleted (purple) cells.

To determine whether the NSL complex physically interacts with the NURF complex, we immunoprecipitated (IP) endogenous NSL complex members (NSL1, MBD-R2, MCRS2) from nuclear S2 cell extracts using polyclonal antibodies. IP experiments successfully enriched for the respective NSL proteins, the NSL complex members (NSL1, NSL3, MCRS2, MBD-R2 and MOF) (Figure 2B) and all four members of the NURF complex (NURF301, ISWI, NURF38 and p55) (Alkhatib and Landry, 2011), albeit substoichiometrically (Figure 2B). To validate these results, we performed IP experiments with an anti-FLAG antibody in cell lines expressing NSL2- FLAG or MBD-R2-FLAG, both of which co-IPed endogenous NSL complex members as well as the endogenous NURF complex members (NURF301, NURF38 and ISWI), but not INO80 (Figure S2C). We could specifically IP endogenous NSL proteins using antibodies raised against NURF301, and ISWI, but not against INO80 and CHD3 (Figure 2C). IP of the chromatin remodeler INO80 exclusively pulls down MCRS2 (Figure 2C). Homologues of MCRS2 and INO80 have been reported to form a complex in mammals (Cai et al., 2006; Chen et al., 2011), which is distinct from the MCRS2-NSL complex (Cai et al., 2010). Consistently, recombinant NURF complex and full length NSL1 but not MCRS2 and MSL3 interact in *in vitro* pulldown assay (Figure S2D). Thus, the NSL complex biochemically interacts with the NURF complex.

To address the relevance of these findings *in vivo*, we assayed the genetic interactions between *nsl1* and the *Nurf301*, *Iswi,* and *Nurf38* members of the NURF complex. Heterozygous *nsl1* mutants show reduced NSL1 activity (Yu et al., 2010) and are 100% viable (Figure S2D, blue bar, Figure S2E). We tested the ability of heterozygous *nsl1* loss-of-function alleles, *nsl1*^*J2E5*^ *(Spradling et al., 1999)* and *nsl1*^*e(nos)1*^ (Yu et al., 2010), to modify the *Nsl3*, *Nurf38*, *Nurf301*, *Iswi* and *Mi-2* RNAi-mediated partial silencing and lethality. We observed a strong negative genetic interaction between *nsl1* and *Nsl3,* as expected for members of the same complex. An equally strong negative interaction was scored between *nsl1* and all tested NURF complex members, whereas no genetic interaction was seen between *nsl1* and *Mi-2*. These findings indicate that NSL1 and NURF interact *in vivo* in the same or parallel converging pathways.

We sought to determine whether this interaction is required only for a specific subset of genes or is a general mechanism. MNase-ChIP-seq experiments against NSL1 and NURF301 revealed that 21,138 (91%) of the 23,194 NSL1 binding sites were also bound by NURF301. Conversely, 66% of the 32,232 NURF301 binding sites were co-occupied by NSL1, and 9,004 (64%) of these co-bound sites overlapped with 14,081 annotated TSSs in the *Drosophila* genome, while only 605 (4%) and 2,182 (15%) were bound by either NSL1 or NURF301, respectively (Figure S2F-2G). Next, we used a CAGE based approach (MAPCap) (see Methods) to map dominant TSSs at single base pair resolution for each gene and sorted the genes by the distance to its closest upstream antisense TSS, within 2,000 base pairs (Figure 2E, Figure S6A). The nucleosomes aligned closely with both the sense and antisense TSS, and both the sense and antisense TSS exhibited extensive co-binding of NSL1 and NURF301. NSL1 and NURF301 co-localized at the NDR upstream of the +1 nucleosome, which is most affected upon NSL1 KD. When we overlaid the signal with TSS positions, the two proteins bind in close proximity to the TSS (Figure 2E). Similar results were obtained when ChIP-seq data for NURF301, ISWI, ACF1 and Mi-2 were analysed (Contrino et al., 2012; Feller et al., 2012). The NSL complex binding sites coincide extensively with the NURF complex but not ACF1 (in the ACF-ISWI complex) or Mi-2 (Figure S3A-C).

To understand the epistatic relationship between NSL and NURF complexes, we performed ChIP, followed by qPCR assays at selected target promoters under either NSL1 or NURF301 KD conditions. KD of NSL1 compromised NURF301 binding to these promoters, while KD of NURF301 left NSL1 binding unchanged (Figure 2F), indicating that the NSL complex acts upstream of NURF complex recruitment. To validate this result, we utilized flies carrying a lexO- luciferase reporter transgene and another transgene expressing lexA-NSL3. Tethering of NSL3 led to ectopic recruitment of NSL1, as well as NURF301 (Figure 2G).

To determine whether recruitment of the NURF complex could explain the defects in nucleosomal organization observed upon NSL1 KD, we depleted NSL1, and the remodelers NURF301, and INO80 in S2 cells (Table S1). MNase-seq experiments revealed that nucleosomes displayed a similar shift towards the TSS upon NURF301 KD (Figure 2H, Figure S4A)(Kwon et al., 2016), while depletion of INO80 led to a shift of nucleosomes away from the TSS. It has been reported that nucleosomes at promoters display differential sensitivity to MNase digestion in *Drosophila* (Chereji et al., 2016). To test if our results are robust in different digestion conditions, we performed our MNase-seq with different various MNase concentrations. Indeed, we can obtain the same conclusion in all digestion conditions (Figure S4B). Interestingly, only NSL1 KD led to a change in nucleosome occupancies, indicating that the NSL complex plays an additional role in maintaining the nucleosome pattern, consistent with the NSL complex being upstream of NURF complex recruitment. Thus firstly, the NSL complex is important for maintaining nucleosome occupancy at +1 nucleosomes and secondly it recruits the NURF complex to position the nucleosomes at active TSSs in the *Drosophila* genome.

### The NSL complex targets TATA-less promoters by recognizing AT content

Since NSL complex appeared upstream of NURF recruitment, we next addressed how NSL complex recognizes target promoters. Previous genomewide correlations suggested association of DRE sequence with NSL target sites (Feller et al., 2012; Lam et al., 2012). However, whether this or other elements could be specifically targeted by the NSL complex remained unknown. For this purpose, we performed DNA immunoprecipitation by isolating and shearing *Drosophila* genomic DNA and incubating it with recombinant NSL1, NSL3 and MCRS2 (Figure S5A). The bound DNA was subsequently purified and sequenced. Unexpectedly, we observed that NSL3, but not NSL1, MCRS2 or GFP (control), binds to specific regions in the genome, characterized by high AT-content (Figure 3A, Figure S5B). We observed that the AT-rich sequences overlap extensively with NSL3 binding *in vivo* (Figure 3B). Interestingly, AT-rich sequences were highly enriched on promoters where TATA and other core promoter motifs were absent (Figure 3B). We calculated the partial correlation coefficient of all 1,024 5mers and found that AT content, rather than a specific motif, correlates best with NSL3 binding. Next, we used our MAPCap TSS as a reference to test whether the AT-content is able to predict NSL targeting. To this end, we used the AT content of 61 29-base pair windows around the MAPCap-based TSSs as predictors for NSL3 *in vivo* binding. We partitioned the genome into training sets and test sets, then trained our logistic regression model with the genes in the training set, and applied the model to predict NSL3 binding on genes in the test sets (ten-fold cross-validation). In this setting, the model correctly predicted 79% of true NSL3 targets and 76% of the non-NSL targets. The high AT content predicts the *in vivo* binding site of NSL3 for bins upstream of MAPCap TSSs, on the other hand, in case of bins overlapped with the +1 nucleosome position, high AT content correlates with lack of NSL3 binding, suggesting that the local chromatin environment also plays a role in directing NSL3 binding (Figure 3C). Furthermore, our analysis revealed that AT-rich sequence is a better predictor than DRE or other 5mer motifs (Figure 3D, 3E). This result suggests that NSL3 recognizes AT rich sequences in the genome and NSL3 binding is further refined by local chromatin context to the restricted locations on housekeeping promoters that lack canonical motifs such as TATA box or Inr.

**Figure 3.**
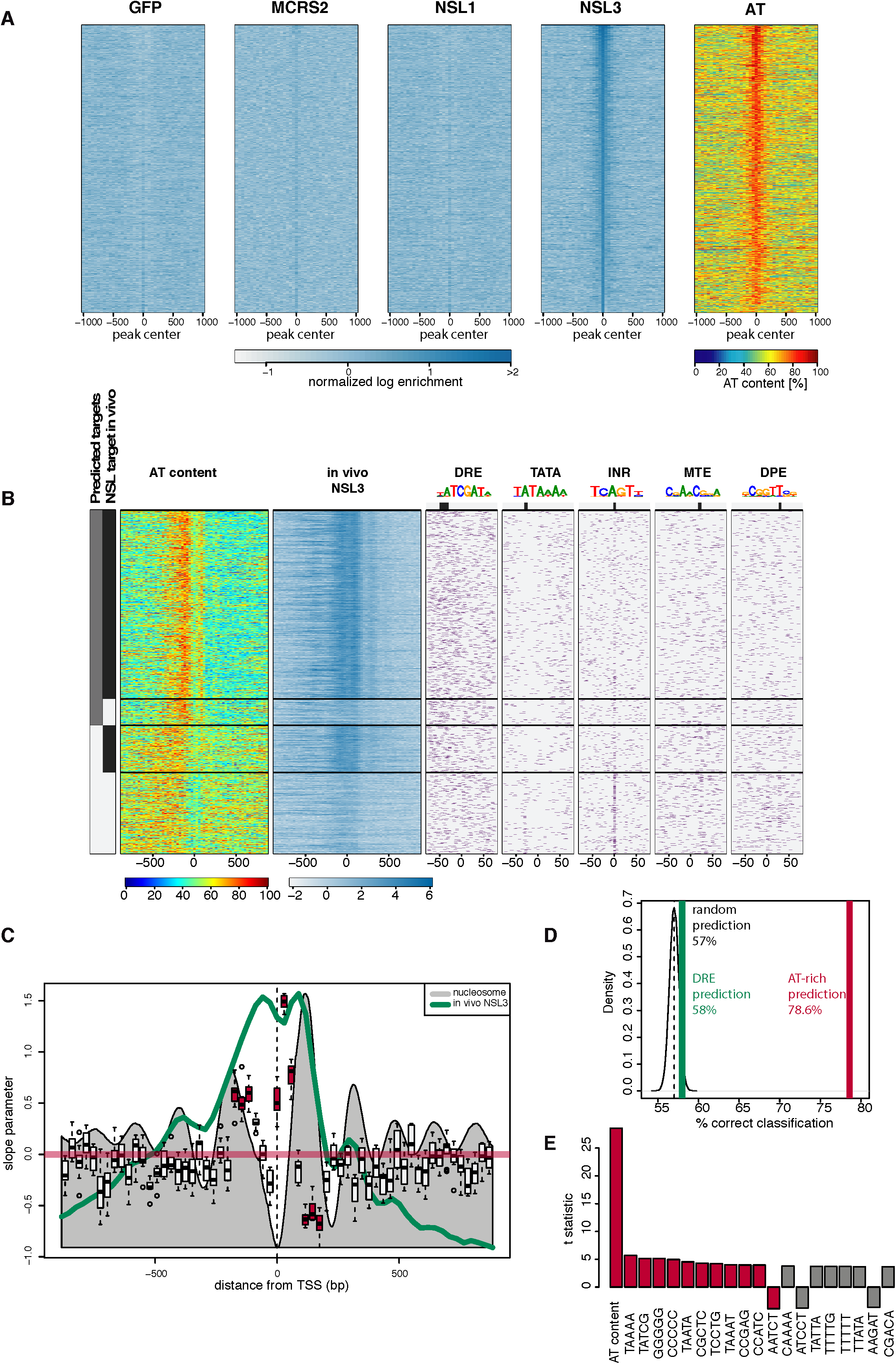
The NSL complex targets TATA-less promoters by recognizing AT content. (A) Heatmaps showing the normalized *in vitro* DNA binding signal of GFP, MCRS2, NSL1 and NSL3 (left to right, blue) using *Drosophila* genomic DNA. The heatmaps are centered around the *in vitro* binding peak center and ordered by binding intensity. AT contents of the same regions are displayed on the rightmost heatmap (red). (B) (leftmost) Bars showing predicted NSL3 binding from AT content (grey) and *in vivo* NSL3 binding as detected by ChIP-seq experiment (black). The first heatmap depicts AT content, while the second heatmap shows *in vivo* NSL3 binding. The following heatmaps on the right show occurrence of PWM hits for core promoter motifs DRE, TATA, INR, MTE and DPE respectively. For each motif a sequence logo and its preferred location is indicated. The genes are clustered into 4 groups along the y-axis: (i) genes predicted to be bound and are bound *in vivo*; (ii) genes predicted to be bound but are not bound *in vivo*; (iii) genes predicted not to be bound but are bound *in vivo* and (iv) genes predicted not to be bound and are not bound *in vivo*. (C) Boxplot showing 29 bp bins which are significantly contributing to prediction of NSL3 binding. The x axis denotes the position with respect to the MapCap TSSs. The y-axis denote the slope parameter values obtained during the 10-fold cross validation to predict NSL3 binding. The grey filled wiggle line denotes the nucleosome signal. The green line denotes the NSL3 *in vivo* binding. The red horizontal line denotes a slope of zero. Red boxes denote bins which successfully predict behaviour of NSL3 binding (t-test, p-value < 0.05). (D) *In vivo* NSL3 binding is predicted using random sequences (black), DRE (green) and AT-rich sequences (red), using the method described in (C). The percentage of sequences making a correct prediction are indicated on the x-axis. (E) Bar charts showing t-statistics representing partial correlation of the indicated elements to *in vitro* binding of NSL3. The AT content (percentage against the log enrichments for all bins) and frequencies of all 1,024 possible 5mers are calculated for 200 bp bins. The *in vitro* NSL3 log enrichment was linearly regressed against the AT content and the frequencies of all possible 5mers used to calculate the partial correlation coefficient for the 5mers.

### The NSL complex is required for transcription start site selection

Consistent with our data, KD of NSL1 should have a strong effect on gene expression and this is indeed what we observed in RNA-seq experiments. The analysis revealed that 5,225 (53%) of the 9,850 genes for which we could detect expression in either control or NSL1 KD were significantly down-regulated at an FDR of 10%, while only 774 (8%) were significantly up-regulated and 3,851 (39%) were not significantly different (Table S2, Figure S6C-D). When considering the log_2_ fold change, 7643 genes were down-regulated (Figure 4A), indicating that the NSL complex is required for transcription of the vast majority of active genes. Notably, there was an overrepresentation of nuclear and mtDNA encoded mitochondrial proteins among the genes that were most down-regulated (Figure 4A). The mammalian NSL complex has been reported to be required for the expression of respiratory genes from both nuclear and mtDNA (Chatterjee et al., 2016). To further investigate the effects on transcription specific to the promoter regions where nucleosome shift occurs, we employed MAPCap analysis in wild type and NSL1-depleted cells (Table S3, Figure S6B). 4,020 (64%) of the 6,281 dominant MAPCap TSSs were significantly down-regulated at a FDR of 10%, while only 113 (2%) were significantly up-regulated and 2,148 (34%) were not significantly changed. Clustering analysis revealed that the TSS expression and nucleosome changes correlate with NSL1 and NURF301 binding (Figure S6C) indicating that changes in the nucleosomal organization are correlated with the down-regulation of TSSs in a NSL1- and NURF301-dependent manner. Promoters targeted by NSL1 and NURF301 show a broader TSS pattern than non-bound ones (Figure 4B), which correlates with the absence of core promoter motifs (Schor et al., 2017) that are predominantly found in non-bound TSSs (Figure S6C).

**Figure 4.**
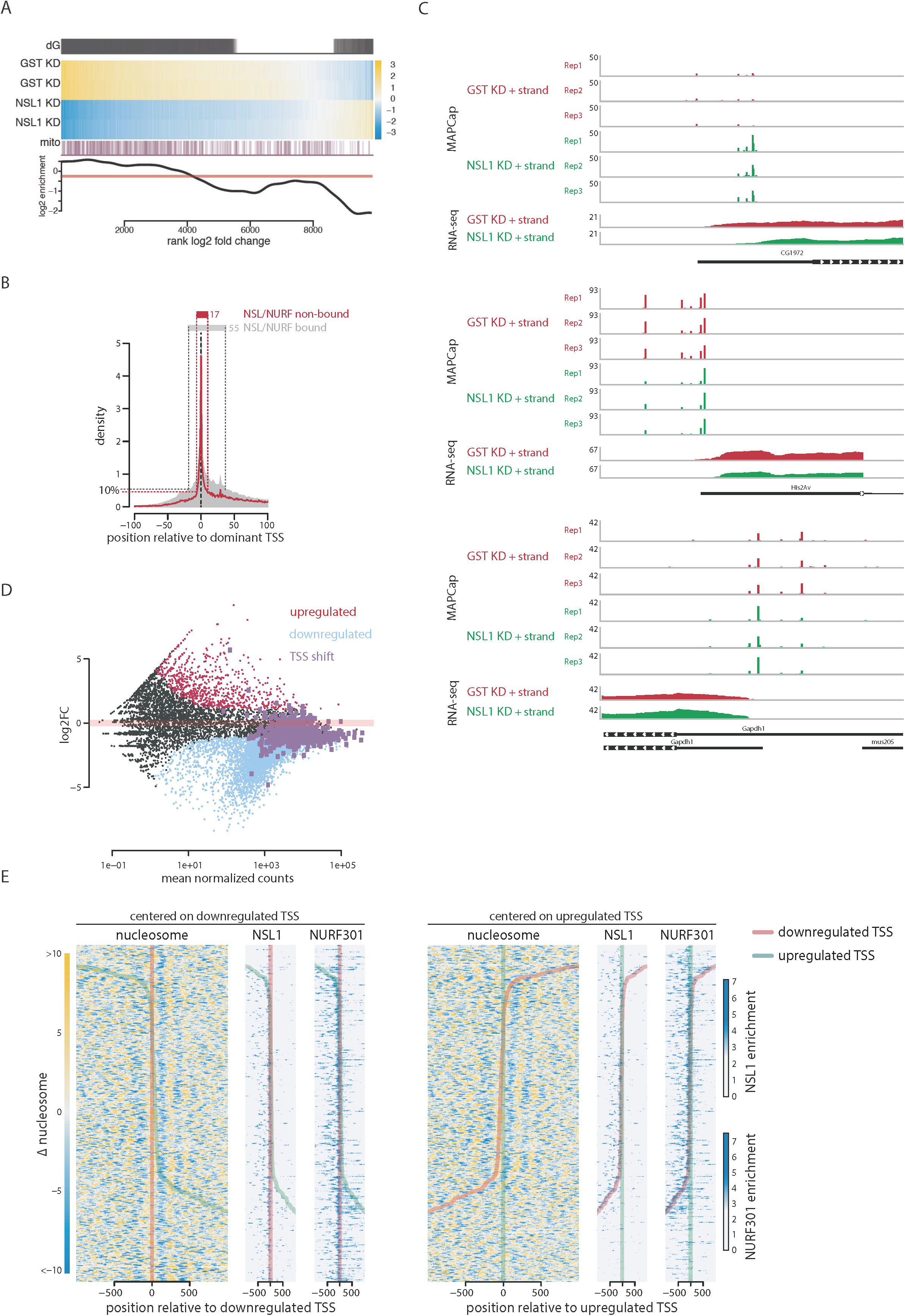
The NSL complex is required for transcription start site selection. (A) (dG) Differentially expressed genes as defined by DESeq2. (heatmap) Gene expression changes in control (GST) and NSL1-depleted cells, yellow and blue color indicate up- and down-regulation respectively, compared to the average of all samples. (mito) Mitochondrial genes are marked as purple lines. The log_2_ enrichment of mitochondrial genes over the transcriptome as background (bottom). (B) The TSS signal obtained from MAPCap is plotted for NSL-bound (grey area) and non-NSL- bound (red line) genes. Signals for each genes are centered around the dominant TSS. The 10 percentile of signal accumulates within 17 bp from the dominant TSS for the NSL-non-bound genes, while the 10 percentile is spread across 55 pb for the NSL-bound genes. (C) Representative examples showing the MAPCap and RNA-seq data for control (GST, red tracks) and NSL1 KD (green tracks). (D) MA plot showing the changes in gene expression upon NSL1 depletion. Light blue color denotes downregulated genes while red color denotes upregulated genes. Genes displaying shifts in TSS selection upon NSL1 KD are marked in purple. (E) Heatmaps showing change in nucleosome occupancy for genes displaying TSS shift in NSL1 KD, with yellow indicating an increase and blue a decrease in nucleosome signal. For each TSS shift event, one TSS is defined as down-regulated (red line in both heatmaps) while another TSS is labeled as up-regulated (green line) in the same gene. Left heatmaps are centered at the down-regulated TSS (red). Heatmaps on the right are centered at the up-regulated TSS (green). NSL1 and NURF301 ChIP-seq signal is shown for these 418 genes.

If the TSS(s) are selected by the position of the +1 nucleosome, a delocalized +1 nucleosome may influence TSS firing and selection. We noticed cases where TSS preference changes upon NSL1 KD (Figure 4C). To identify genes that change their TSS preference upon NSL1 KD, we devised a statistical analysis (see Methods) that identified 418 genes with a significant change in TSS usage at a FDR of 5%. Some of these genes alter their promoter usage, and 251 (60%) exhibit changes in the TSS usage within a window of 200 base pairs, often with one TSS being favored while another is repressed. We asked how this change in TSS usage alters overall gene expression and found that 6 (1%) genes were up-regulated, 249 (60%) genes remained unchanged, and 162 (39%) genes were down-regulated (Figure 4D). Overall expression of most of these genes remained unchanged, indicating that an alternate promoter or TSS compensates for the depressed NSL1-regulated TSSs. These genes were bound by both NSL1 and NURF301 (Figure 4E). We therefore asked whether this change in TSS preference could be a direct consequence of the changes in the nucleosomal pattern upon NSL1 KD. For each of the 418 genes, we identified the TSS that is favored in NSL1 KD as an up-regulated TSS while the TSS which showed reduced usage was marked as a down-regulated TSS. The down-regulated TSSs aligned with a local increase in nucleosome occupancy in their NDRs (Figure 4E), indicating that the disruption of the canonical nucleosomal organization leads to a down-regulation of certain TSSs.

The usage of an alternate promoter could have important consequences for the resulting mRNA. For example, a shift in TSS usage within a promoter can result in changes in 5’ UTR length which may affect post-transcriptional regulation of the resulting mRNA (Hinnebusch et al., 2016; Leppek et al., 2017). We compared the RNA-seq coverage in these regions from control and NSL1 KD samples, focusing our analysis on genes that had RNA-seq coverage within a window of +/- 200 base pairs around the TSS. In cases when a downstream TSS was up-regulated, we clearly observed a reduction of RNA-seq coverage at the beginning of the 5’UTR, indicating shorter transcripts. Likewise, when an upstream TSS was up-regulated, we observed more RNA-seq reads upstream of the 5’UTR and hence a longer transcript (Figure 5A). Consistently, comparison of NSL1-mediated changes in TSS usage with ribo-seq data (Dunn et al., 2013) revealed that TSS shifts upon loss of NSL1 would impact the 5’ UTR as well as the translated products in cases where a TSS shift occurred downstream (Figure 5B, Figure S6E). Thus, the NSL complex can influence TSS choice through canonical nucleosomal organization.

**Figure 5.**
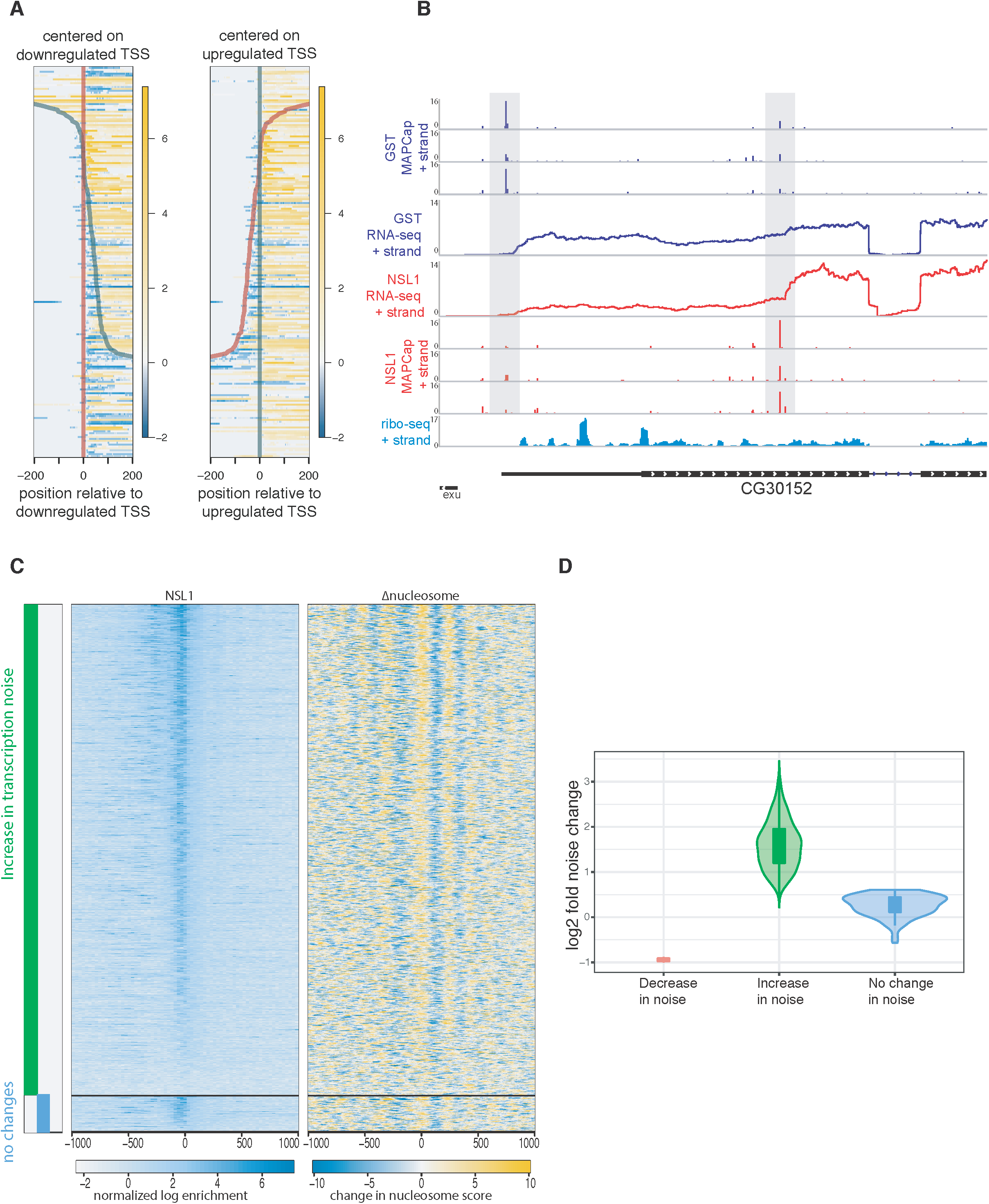
Transcription noise increases in the absence of the NSL complex. (A) Differential signal in RNA-seq comparing control and NSL1 KD at 5’UTR regions. Heatmaps are centered at the down-regulated (left) TSS and up-regulated TSS (right) as in Figure 4C. Yellow denotes an increase in RNA-seq read counts, while blue denotes a decrease in RNA- seq read counts and thus represent changes in the length of 5’UTR. (B) Representative example showing MAPCap as well as RNA-seq data for control and NSL1 KD samples. Ribo-seq data from wild type cells is provided (bottom). Ribo-seq data shows upstream ORF in the 5’UTR region, where expression is reduced in NSL1 KD due to a shift in TSS selection. (C) (Leftmost) Bars showing genes with increased (green) and unchanged (blue) transcription variability. Differential variability was calculated using the BASiCS_TestDE function. The BASiCS_TestDE function also corrects for changes in gene expression between control and NSL1 KD. (middle and right) Heatmaps showing NSL1 binding and changes in nucleosome occupancies upon NSL1 KD on the same genes. (D)Violin plot showing the change in transcription variability depicted by the green and blue bars in (C).

**Figure 6.**
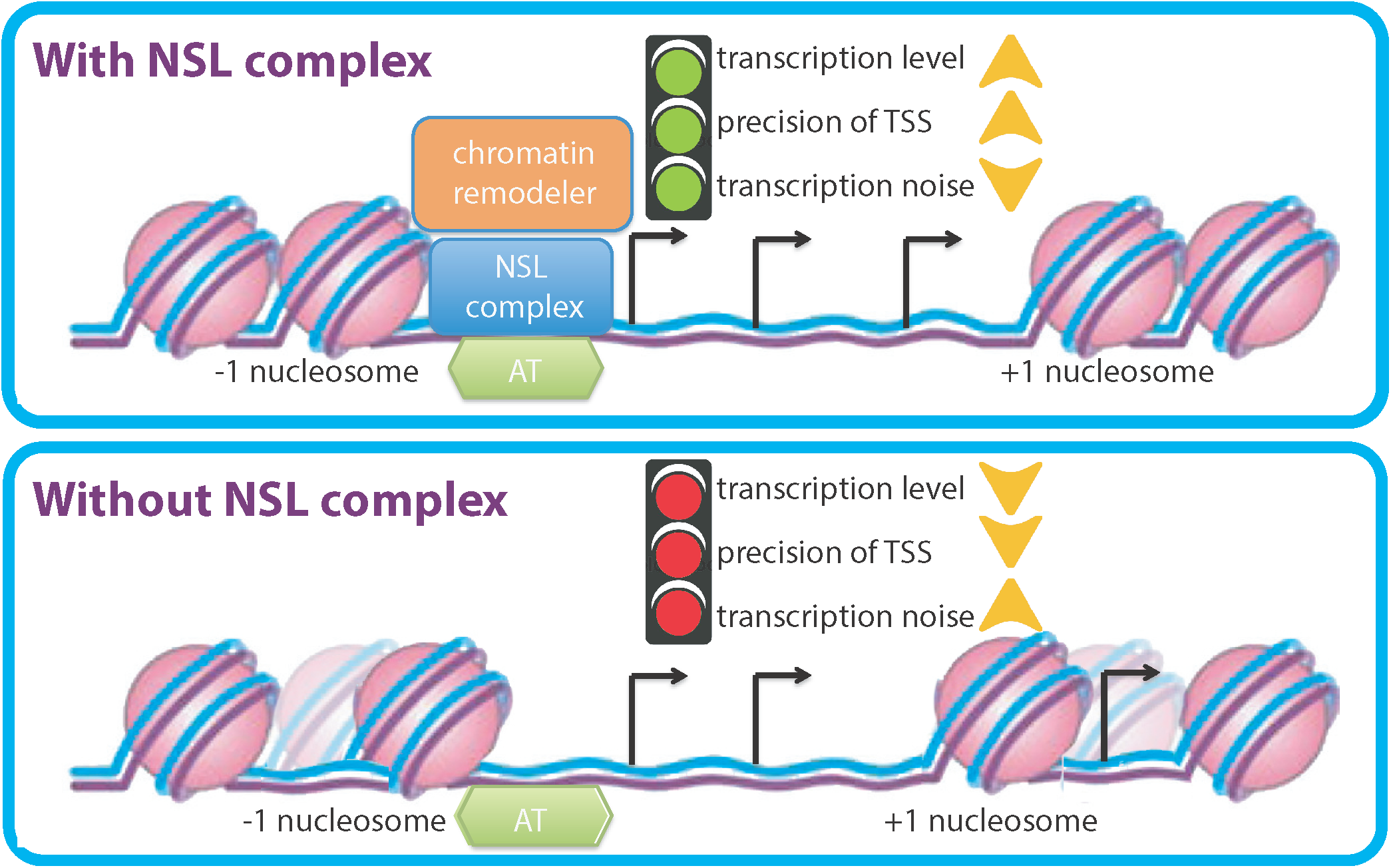
A schematic model. The NSL complex is required for nucleosome positioning at dispersed promoters of housekeeping genes. Genetic and biochemical analysis revealed that the NSL complex recruits NURF chromatin remodeling complex at target promoters. The NSL complex targets AT rich sequences in TATA less promoters. Loss of NSL complex not only leads to transcription downregulation of targets, but in addition affects choice of transcription start sites of highly expressed genes with multiple TSSs. Loss of NSL complex also increases transcription noise at target genes. Thus, NSL complex plays a crucial role in transcription fidelity in the Drosophila genome.

### Transcription noise increases in the absence of the NSL complex

Our results suggest that the NSL complex creates a transcription competent nucleosome organization at its target promoters enabling constitutive expression with concomitant low transcription noise (Eldar and Elowitz, 2010; Lehner, 2010; Munsky et al., 2012; Ravarani et al., 2016; Sanchez et al., 2013; Sharon et al., 2014). Disruption of this transcription competent nucleosome organization at the TSS should lead to a TSS that requires chromatin remodeling prior to transcription such that it cycles between an off and an on state, which increases transcription noise (Eldar and Elowitz, 2010; Munsky et al., 2012). Therefore, we asked whether the disruption of the nucleosome organization by the depletion of NSL1 results in an increase in transcription noise. For this purpose, we performed single-cell RNA-seq on control *Drosophila* S2 cells and cells depleted of NSL1 using the 10x genomics emulsion-based sequencing technology. 1753 cells and 4046 cells were sequenced for NSL-depleted and control samples, respectively. We selected cells with more than 2,000 expressed genes and focused on a subset of genes that were expressed in more than 50% of the remaining cells. To detect changes in transcription noise we used the BASiCS approach (Vallejos et al., 2015). Since transcription noise is influenced by transcription levels, we focused on 1,008 genes which were not differentially expressed upon NSL1 KD and identified changes in transcription noise at an expected false discovery rate of 10%. We observed increased transcription noise in most of these genes (Figure 5C, 5D, Figure S7). Furthermore, the majority of genes with increased transcription noise were NSL1 targets and displayed disrupted nucleosome patterning upon NSL1 KD (Figure 5C). Thus, our data suggest that the NSL complex is involved in suppressing transcriptional noise at target gene loci.

## Discussion

In the current study, we set out to understand how transcription initiation is regulated on NSL- bound promoters, which are typically TATA-less housekeeping promoters with dispersed TSS pattern. We uncover here that the NSL complex is required for maintaining the positioning of the +1 nucleosome at NSL-bound gene promoters, which is not only pivotal for effective transcription but also for TSS fidelity in *Drosophila*.

We find here that the NSL complex recruits the NURF chromatin remodeling complex to NSL target promoters. In mammals, BPTF (NURF301 homolog) binds to promoter-associated H3K4me2/3 and H4K16ac via its PHD and bromodomain respectively, and the interaction is important for the recruitment of the NURF complex (Ruthenburg et al., 2011; Ruthenburg et al., 2007). In *Drosophila*, two isoforms of NURF301 exist. A longer isoform which encodes the C-terminal PHD and bromodomain, as well as a shorter isoform that lacks the two domains. The shorter isoform thus is unable to bind H3K4me3 or H4K16ac, but is nevertheless sufficient to target the NURF complex to the majority of genes (Kwon et al., 2009), which then acts upon the +1 nucleosomes to properly position them. Our data suggest a plausible explanation on how the NURF complex may be recruited by transcription factors such as the NSL complex, in addition to the histone marks. It is thus conceivable that the NURF complex interacts with both the NSL complex and the histone marks for accurate targeting to gene promoters.

Transcription stochasticity is known to play an instrumental role in processes such as cell differentiation, as well as cellular homeostasis in response to external stimuli (Eldar and Elowitz, 2010; Munsky et al., 2012). TATA-containing promoters are believed to be more noisy and important for fate determination while TATA-less promoters are considered to be less noisy and involved in cellular homeostasis (Shalek et al., 2013). However, it was previously not clear how TATA-less housekeeping genes regulate transcription noise levels. Our work here reveals that the NSL complex plays a role in suppression of transcription noise at housekeeping genes. We show that nucleosome occupancies at NDR increase in the absence of the NSL complex. This increase likely posits that nucleosome occupancy of a certain promoter at a given time is more heterogenous between cells. Consistent with previous reports that demonstrated that DNA sequences with low nucleosomal affinity display lower transcriptional noise (Dadiani et al., 2013; Sharon et al., 2014), our data suggest that the increase in heterogeneity of nucleosome occupancy leads to an increase in transcriptional noise as observed upon NSL depletion. Moreover, the NSL complex has been shown to facilitate the recruitment of Pol II machinery (Lam et al., 2012). Thus, it is also plausible that the NSL complex changes the dynamics of Pol II binding and initiation at chromatin and thereby represses transcriptional noise at NSL-bound genes. Our results have implication for further understanding how transcription noise can be regulated by transcription factors on distinct types of promoters.

Variants in core promoter DNA sequence affect TSS firing pattern in different *Drosophila* lines (Schor et al., 2017). Yet, it is not understood how these changes in DNA sequence are translated to changes in TSS pattern. Our MAPCap analyses in NSL1 KD cells reveal important insights into the crucial roles that the NSL complex binding and nucleosome positioning plays in ensuring efficient TSS firing and selection. For most genes, TSS firing is compromised as nucleosomes invade the NDR in the absence of the NSL complex. Some genes compensate for the reduced TSS firing by upregulating TSS firing from an alternative promoter of the same gene. This surprising result suggests that neighboring TSSs within one gene could rely on different mechanisms for initiation. It is thus possible that different TSSs may be preferred in different tissues or stress conditions in wild type cells. This TSS preference shift may potentially be regulated by modulating nucleosome positioning or binding of the NSL complex. The mis-regulation in TSS preferences upon NSL1 KD is not without consequence. The TSS shift produces a RNA with altered 5’UTR length and in some cases, an altered 5’ nucleotide. The altered UTR length leads to different numbers of upstream open reading frames (uORFs) and RNA modifications in the UTR while a changed starting nucleotide affects the frequency of m6Am modification. uORFs and RNA modifications have well known effects on RNA stability and translation (Barbosa et al., 2013; Mauer et al., 2017), suggesting significant changes in the cellular proteome in the absence of the NSL complex.

Housekeeping TATA-less promoters are enriched in DNA motifs such as motif 1, 6, 7 and DRE (Ohler et al., 2002). Nevertheless, no single motif is enriched on the majority of housekeeping promoters, raising the questions: is there an equivalent to TBP/TATA box on housekeeping promoters and how these promoters are targeted specifically? Our data uncover that the NSL complex targets these TATA-less promoters by recognizing AT-rich sequences. Remarkably, using AT content and chromatin context, we can correctly predict 79% of true NSL3 *in vivo* targets and 76% of the non-NSL targets. Since the NSL complex binds to the majority of housekeeping promoters, our result shed light on how transcription factors can discern TATA-less promoters from the rest of the genome.

In summary, we show that the NSL complex functions to maintain a prominent promoter nucleosome pattern and subsequently guides TSS selection and suppresses transcription noise on dispersed housekeeping promoters. Our data also reveal that the NSL complex is recruited to the majority of TSSs that lack canonical promoter motifs such as the TATA-box and Inr by binding to AT rich sequences. Taken together, this study provides a plausible explanation to the long-standing questions of how TATA-less promoters are recognized and transcribed. Based on our data, we propose a model whereby the NSL complex acts like a Swiss-army knife/platform to bring together the characteristic chromatin modifying factors that are typically observed at housekeeping genes and required for their proper transcription.

## Acknowledgments

We are grateful to Nicola Iovino and Bilal Sheikh for critical reading of the manuscript and the MPIIE core Deep sequencing facility and bioinformatic department. We thank Erika Pearce for providing access to 10xgenomics technology. We thank Carl Wu and Paul Badenhorst for providing reagents. This work was supported by CRC992, CRC1140 and CRC746 awarded to AA.

## Data Accession

All the genomewide datasets in this manuscript have been deposited in GEO: Accession number: **GSE118726**

Reviewers/Editor access code: **upylqyworribxox**

## Supplementary Information

**Supplementary Figure 1.**
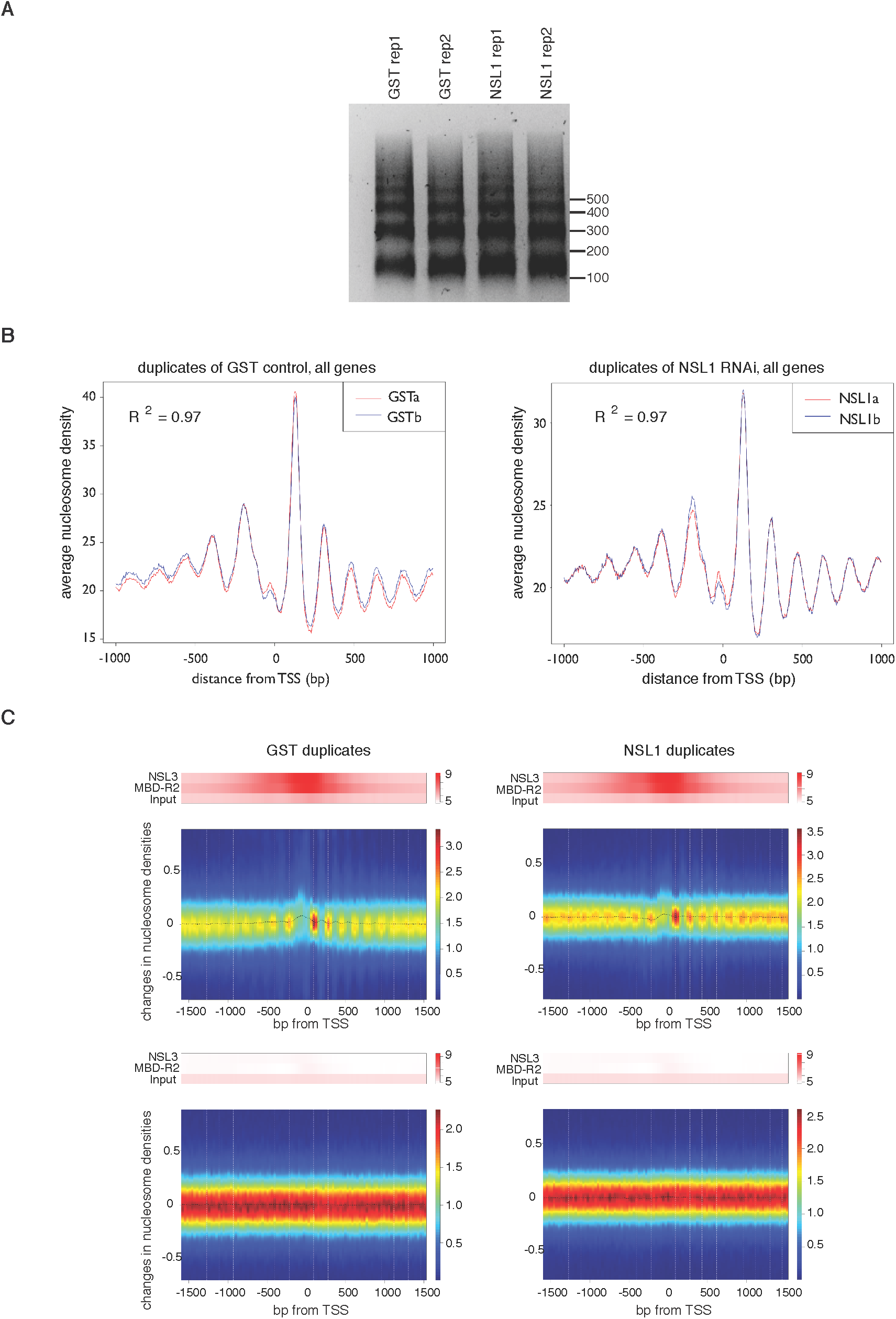
(A) Digestion pattern of samples prepared for MNase-seq experiments. Knockdown experiments using dsRNA against GST (control) NSL1 are done in duplicates. (B) Summary plots showing correlation between duplicates in MNase-seq experiments. R^2^ values are calculated from Spearman correlation coefficient. A region 1 kb around the TSS is shown. (C) Heatmap showing the differences in nucleosome densities between replicates of control (GST; left) and NSL1 KD (right) for active (above) and inactive (below) genes. The y-axis represents changes in nucleosome signals. The color scale bar indicates scatter density. ChIP-seq signals of NSL3 and MBD-R2 are represented in bars above the heatmap to show the binding sites of the proteins. A region 1.5 kb around the TSS is shown.

**Supplementary Figure 2.**
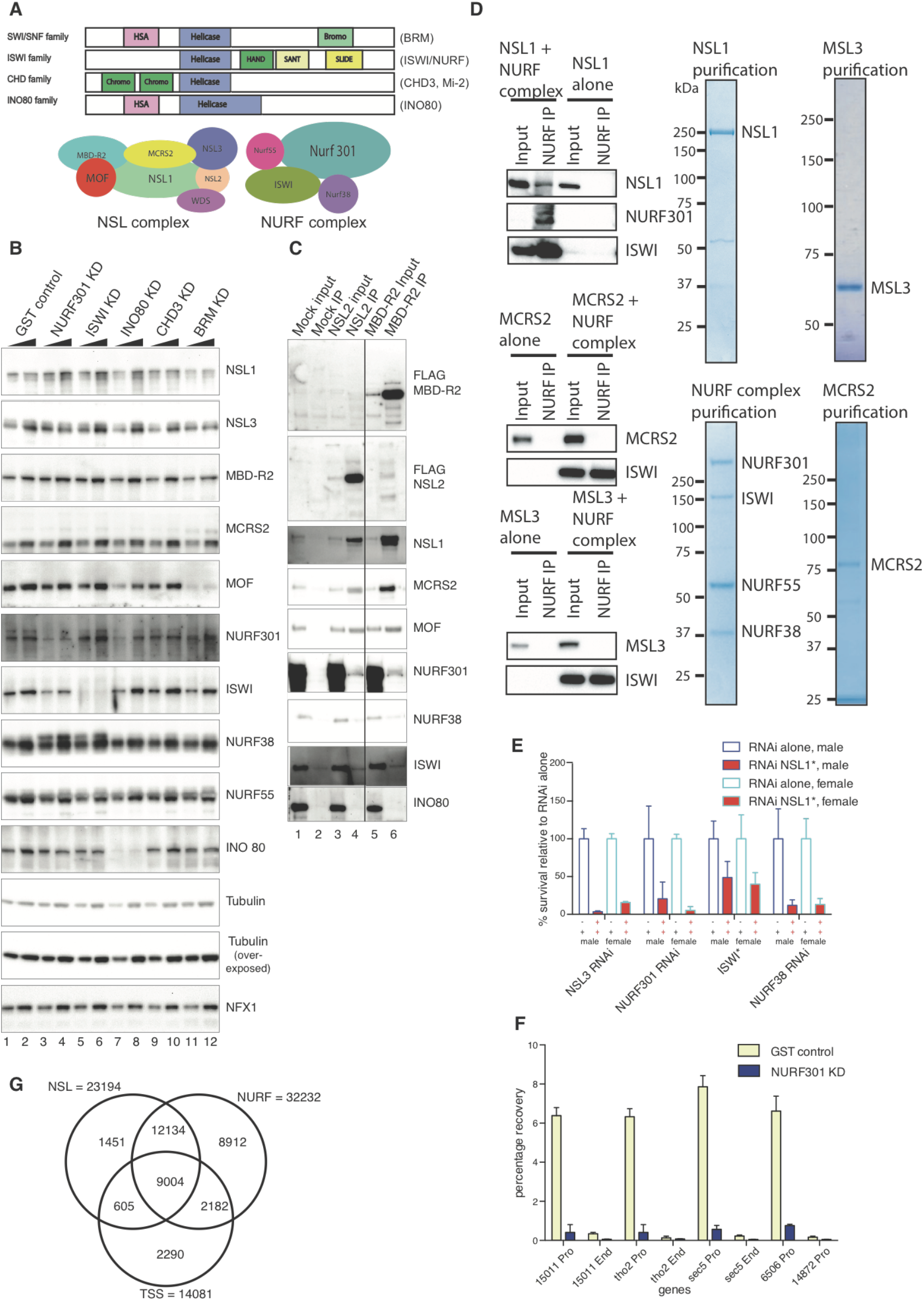
(A) (Top) Schematic representation of 4 families of chromatin remodelers. Proteins of the respective family used in this study are shown on the right. (Bottom) Cartoon representation of core NSL complex and NURF complex members. (B) Western blot showing protein levels of the NSL complex and chromatin remodelers in NURF301, ISWI, INO80 and CHD3 KD. The western blots show that the KD efficiently deplete the respective remodelers. The level of MOF is significantly reduced in the BRM KD. (C) Western blot analysis of immunoprecipitation of exogenous FLAG-tagged NSL2 and MBD-R2 proteins using anti-FLAG peptides antibody. The protein bands are detected with antibodies against the NSL complex members, NURF38, NURF301, ISWI and INO80. (D) Western blot analysis of immunoprecipitation of *in vitro* expressed HA-tagged ISWI using anti-HA peptides antibody. The NURF complex, including HA-ISWI, was incubated with NSL1, MCRS2 (member shared by the INO80 and NSL complex) or MSL3 (member of the MSL/MOF complex). Coomassie protein gels show individual purification of NSL1, MSL3, MCRS2 and NURF complex used in the experiment. KDa denotes molecular weight markers. (E) Genetic interaction between the NSL and the NURF complex as in Figure 2D. NSL1 mutant shows genetic interaction with NURF301, ISWI and NURF38 in both males and females. (F) ChIP-qPCR analysis using NURF301 antibody in wild type and NURF301-depleted S2 cells. The antibody specifically detects NURF301 signal in the wild type but not in the KD samples. Percentage of input recovery is shown on the y-axis. (G) Venn diagram showing overlapping binding sites of NSL1 and NURF301 as detected by ChIP-seq experiments.

**Supplementary Figure 3.**
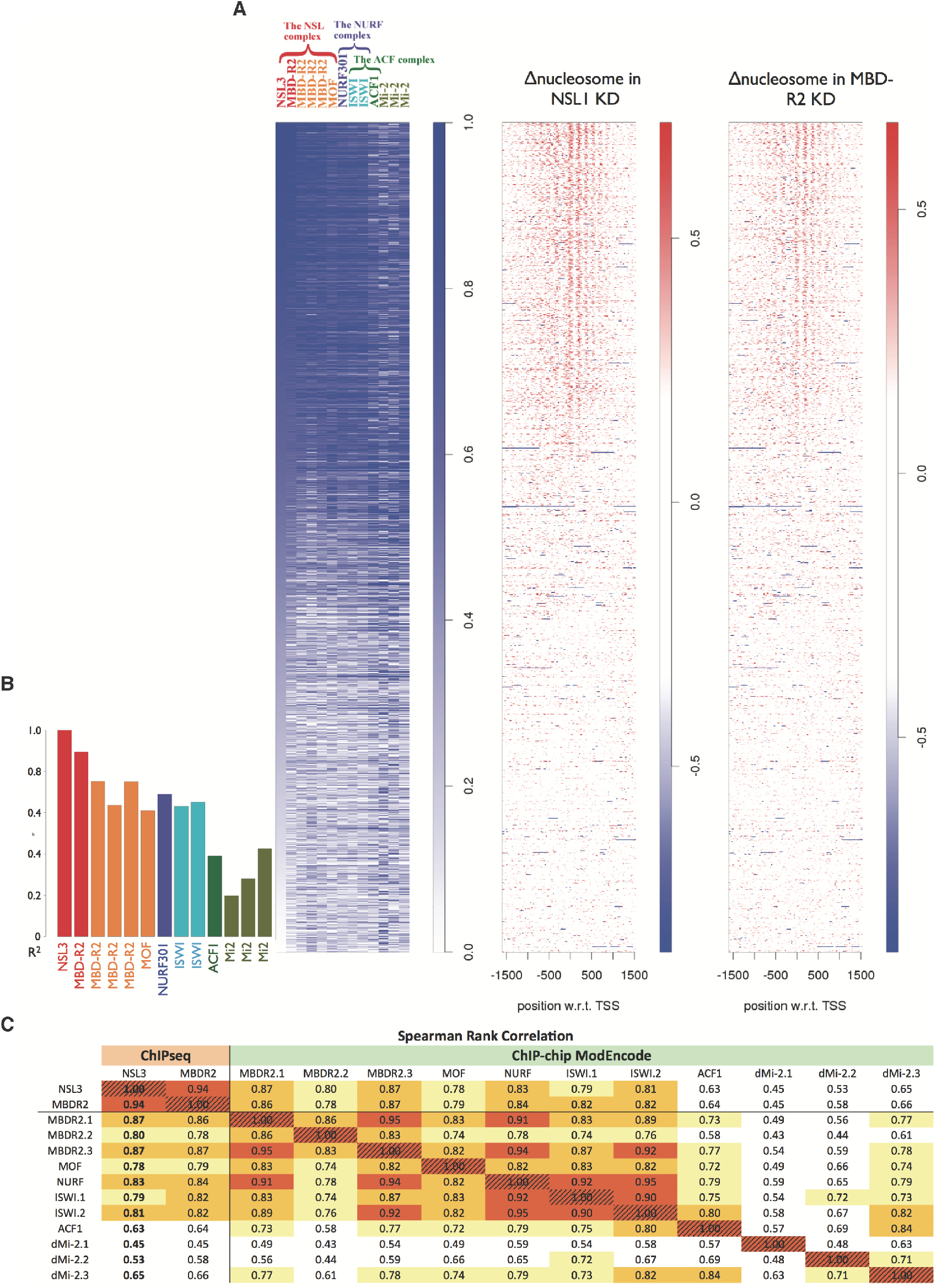
(A) The NSL complex binding sites are more similar to those of the NURF complex than those of ACF1 or Mi-2 (leftmost heatmap) Heatmap showing the promoter (TSS±200 bp) binding signal of (left to right) NSL3 and MBD-R2 (ChIP-seq, (Lam et al., 2012)) as well as MBD-R2 (in triplicates), MOF, NURF301, ISWI (in duplicates), ACF1 and Mi-2 (in triplicates) (ModENCODE). The intensity of the blue color indicates the rank of enrichment signal. (central and rightmost) Heatmaps showing the changes in nucleosome density on the respective genes for the region TSS ± 1.5 kb. The intensity of the red color indicates the degree of nucleosome loss. All heatmaps were in the same gene order arranged according to the NSL3 (ChIP-seq) enrichment. (B) The bar chart displays the spearman correlation coefficients of the NSL3 ranks against ranks in other tracks. There are high correlations between binding of NSL3 and the binding of NURF301 and ISWI and NSL3 binding is less correlated with the ACF1 (also interact with ISWI) or Mi-2. (C) All pairwise Spearman rank correlation coefficients can be found in the table. Data in the first column of the table, where the numbers are in bold, was taken to plot the bar chart mentioned above. The table is color coded: with red (0.9) and orange (0.8) represent highly correlated pairs of profiles, while yellow (0.7) and white (<0.7) represent lowly correlated pairs. All tracks are ChIP-chip data from ModENCODE, unless otherwise stated. These data are publicly available and were retrieved from the ModENCODE project website (www.modencode.org)

**Supplementary Figure 4.**
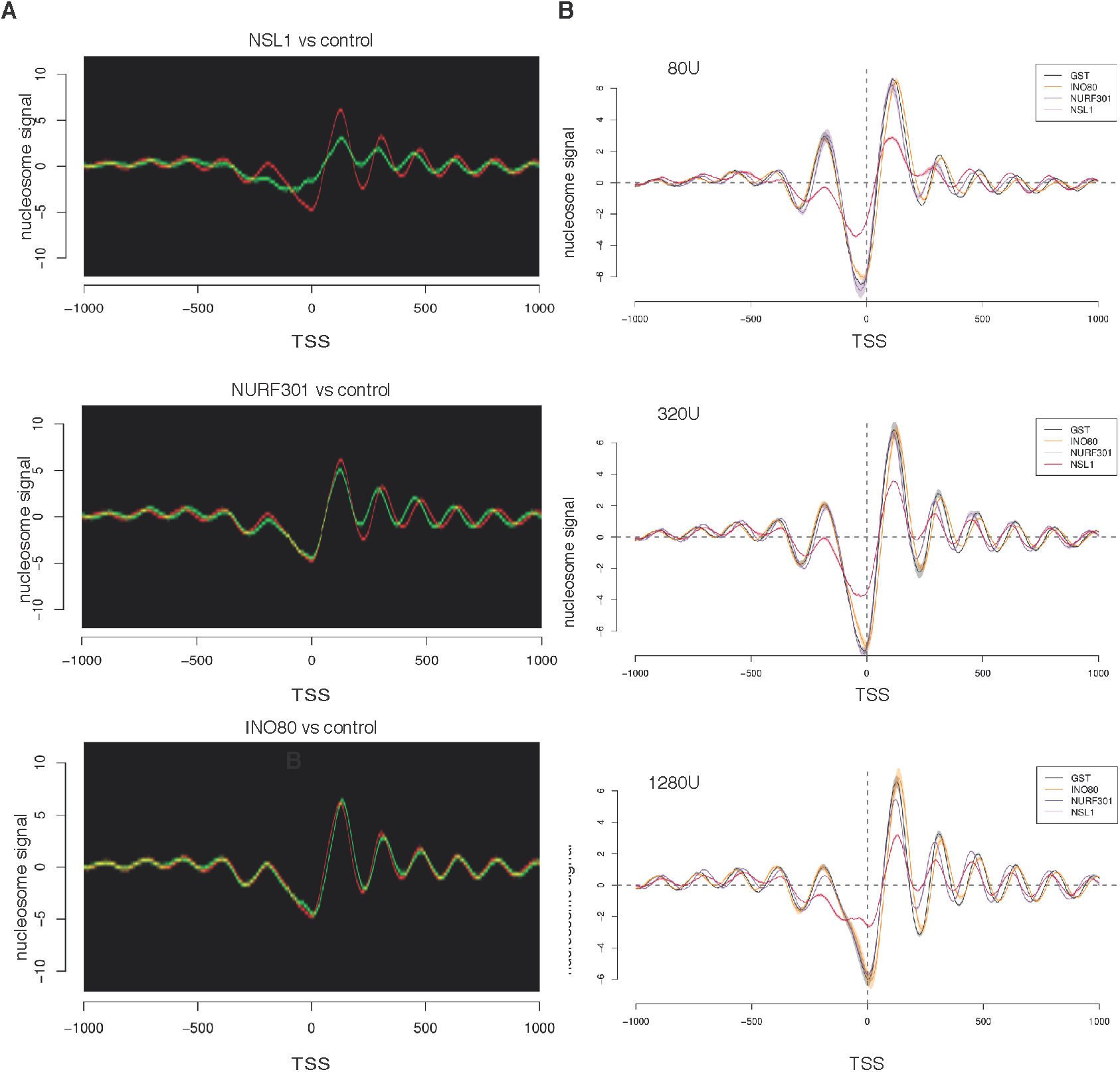
(A) Summary plot showing nucleosome shift in NSL1, INO80 and NURF301 KD. Nucleosome signals of MNase-seq are plotted on the y-axis. Red lines show the control experiments and green lines show the KD experiments. A region 1 kb around the TSS is shown. (B) MNase-seq experiments were reproduced with different degree of MNase digestions. Mnase-seq experiments are repeated with 80, 320 and 1280U of MNase in control, NSL1, NURF301 and INO80 depleted cells. The shaded area around the lines represent the variation between replicates.

**Supplementary Figure 5.**
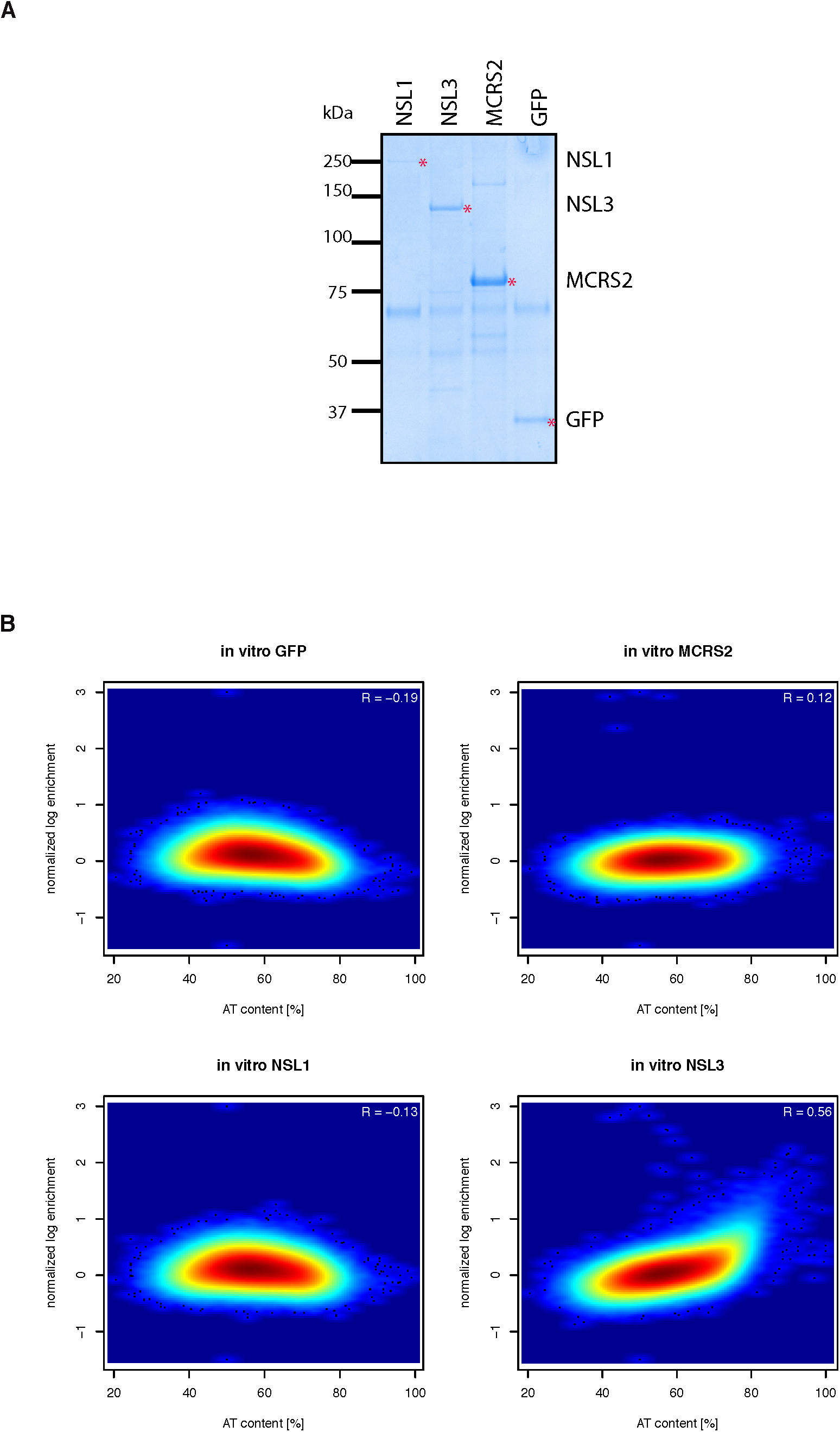
(A) Protein gels showing purification of NSL1, NSL3, MCRS2 and GFP used for the genomic DNA immunoprecipitation experiment. KDa represents molecular weight markers. (B) Scatter plots showing correlation between *in vitro* binding of GFP, MCRS2, NSL1, NSL3 and AT content.

**Supplementary Figure 6.**
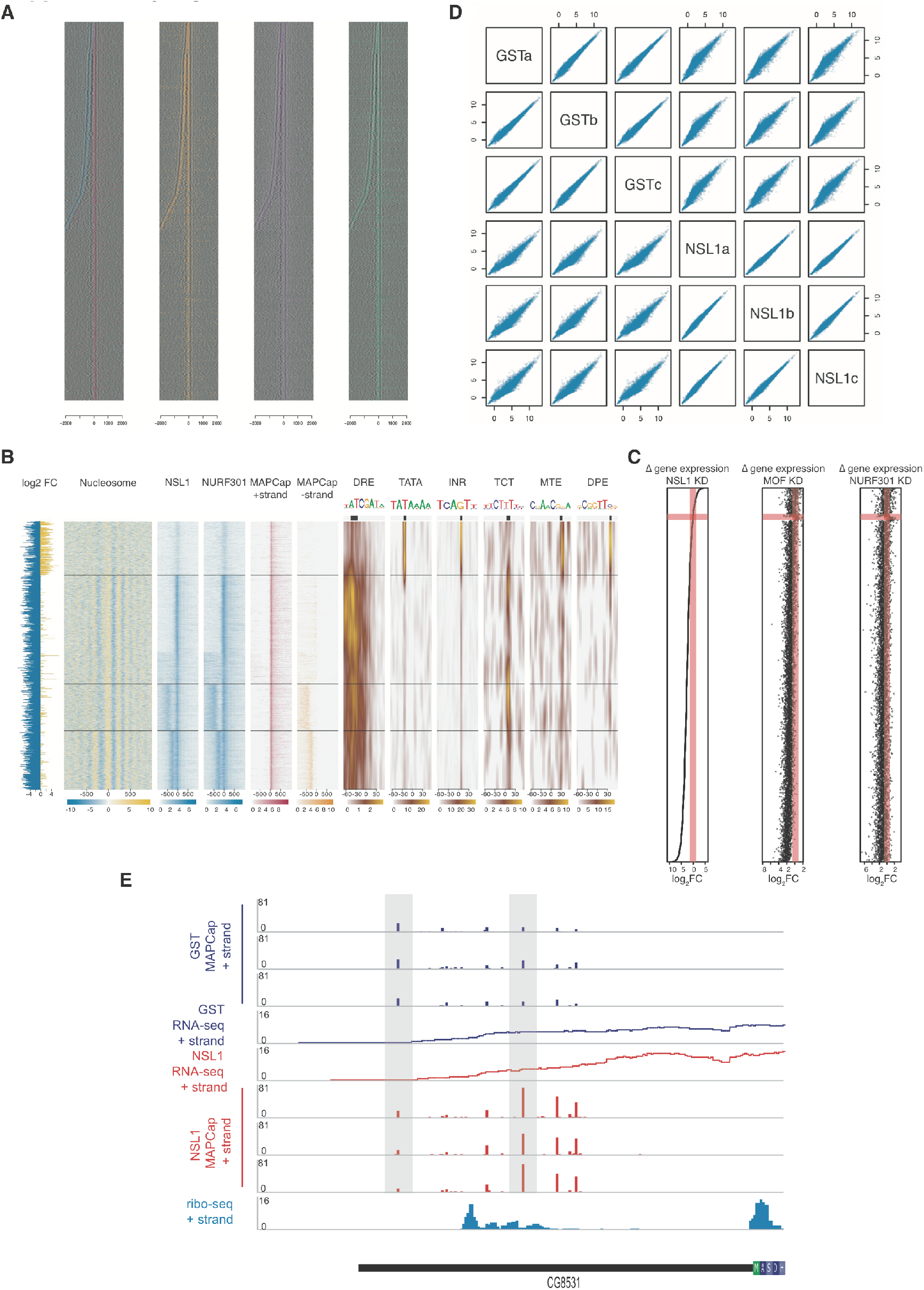
(A) Heatmaps showing the overlap between TSS positions (MAPCap), NSL1 (orange), NURF301 (purple), Pol II (Rpb3, green) and nucleosome pattern (black). (B) Scatter plots showing correlation between triplicates of the MAPCap data. Shown are the normalized and regularized log2 count values as computed by the rlog function of DESeq2. Triplicates of the GST and NSL1 samples show good correlation. (C) Dominant MAPCap TSSs are clustered into five groups using the k-means clustering algorithm with the NSL1 and NURF301 ChIP-seq signal pattern. For these five groups of TSSs, we show (from left to right) tracks showing changes in gene expression upon NSL1 depletion. Blue denotes gene downregulation, yellow denotes gene upregulation. (1^st^ heatmap) changes in nucleosome signal upon NSL1 depletion, yellow indicates gain and blue indicate loss in signal. (2^nd^- 3^rd^ heatmap) ChIP-seq signal of NSL1 and NURF301. (4^th^-5^th^ heatmap) MAPCap data showing the position of TSS on plus (pink) and minus (orange) strands (6^th^ – 11^th^ heatmap) motif enrichment analysis for *Drosophila* core promoter motifs. The color indicates the enrichment. (D) Scatter plots showing changes in gene expression in NSL1, NURF301 or MOF depleted cells. Genes are ordered according to the gene expression changes in NSL1 KD (leftmost). (E) Representative example showing MAPCap, RNA-seq data for control and NSL1 KD samples, and ribo-seq data for wild type cells. Ribo-seq data shows upstream ORF in the 5’UTR region, where expression is reduced in NSL1 KD due to shift in TSS selection.

**Supplementary Figure 7.**
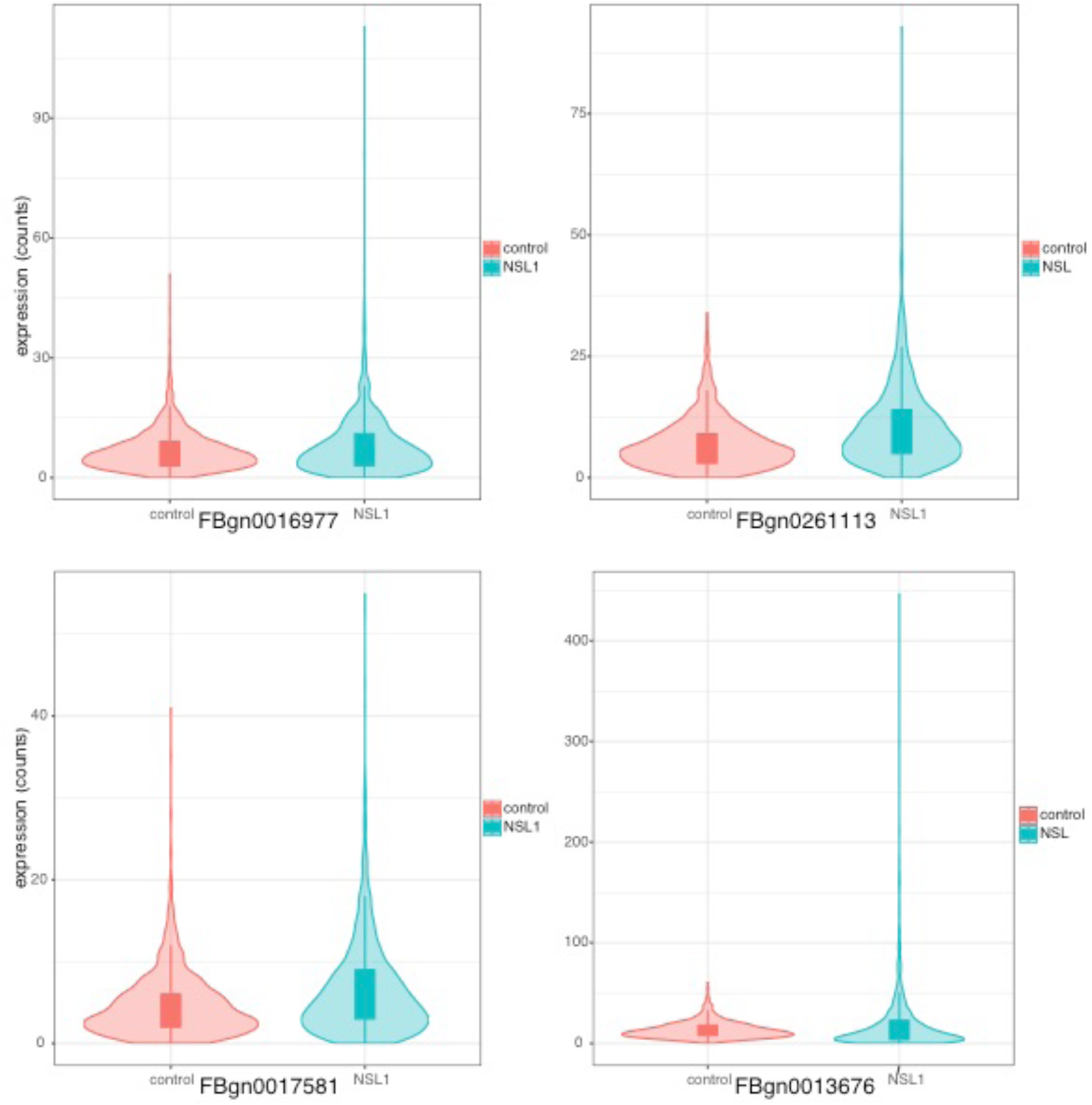
Violin plots showing representative examples of changes in expression noise between control (red) and NSL1 KD (green). The lower and upper hinges of the boxplot correspond to the first and third quartiles while length of the whiskers represents 1.5 times the interquartile range.

**Table S4.**
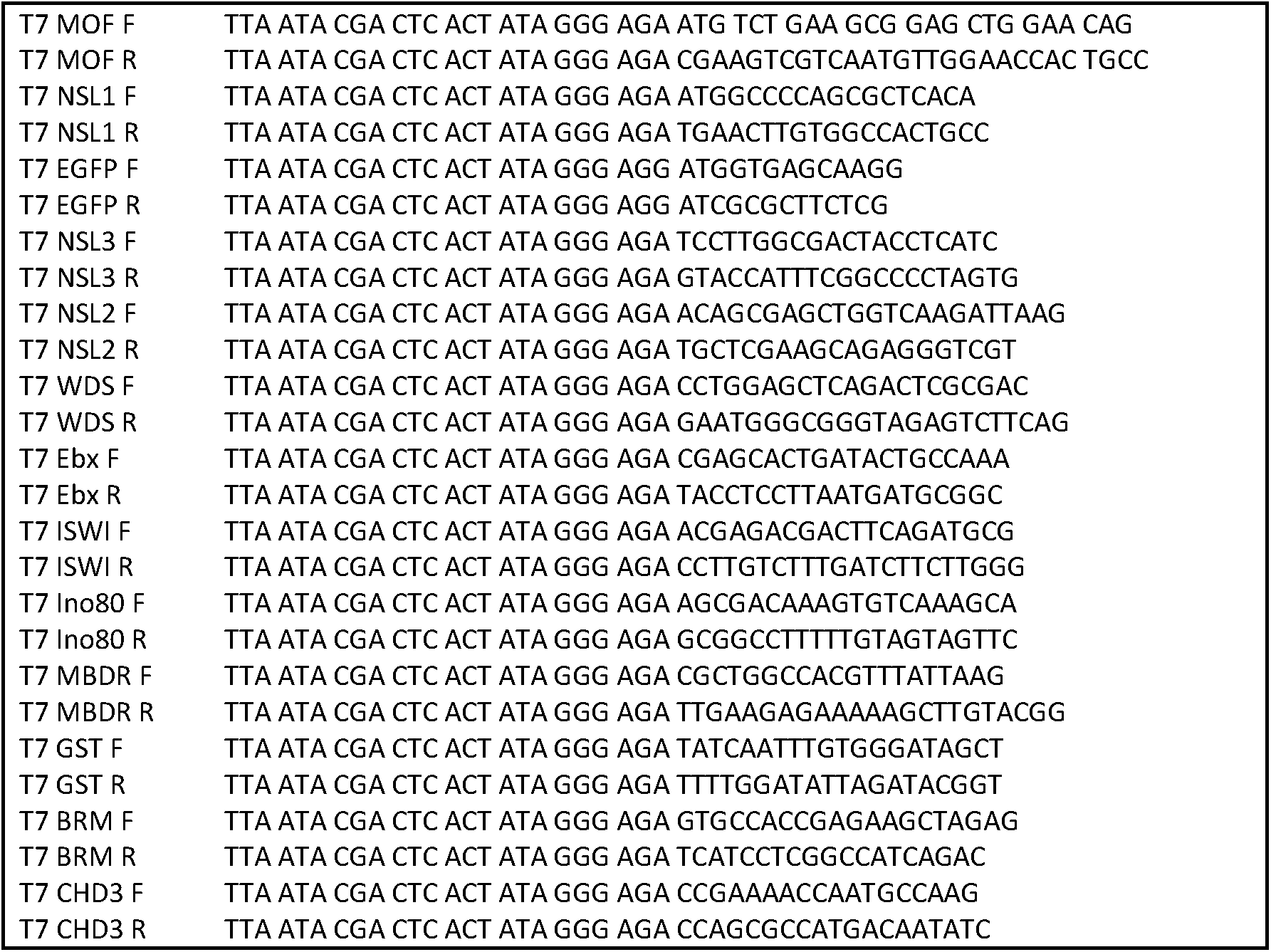
List of primers used for amplification of dsRNA for knockdowns.

**Table S5.**
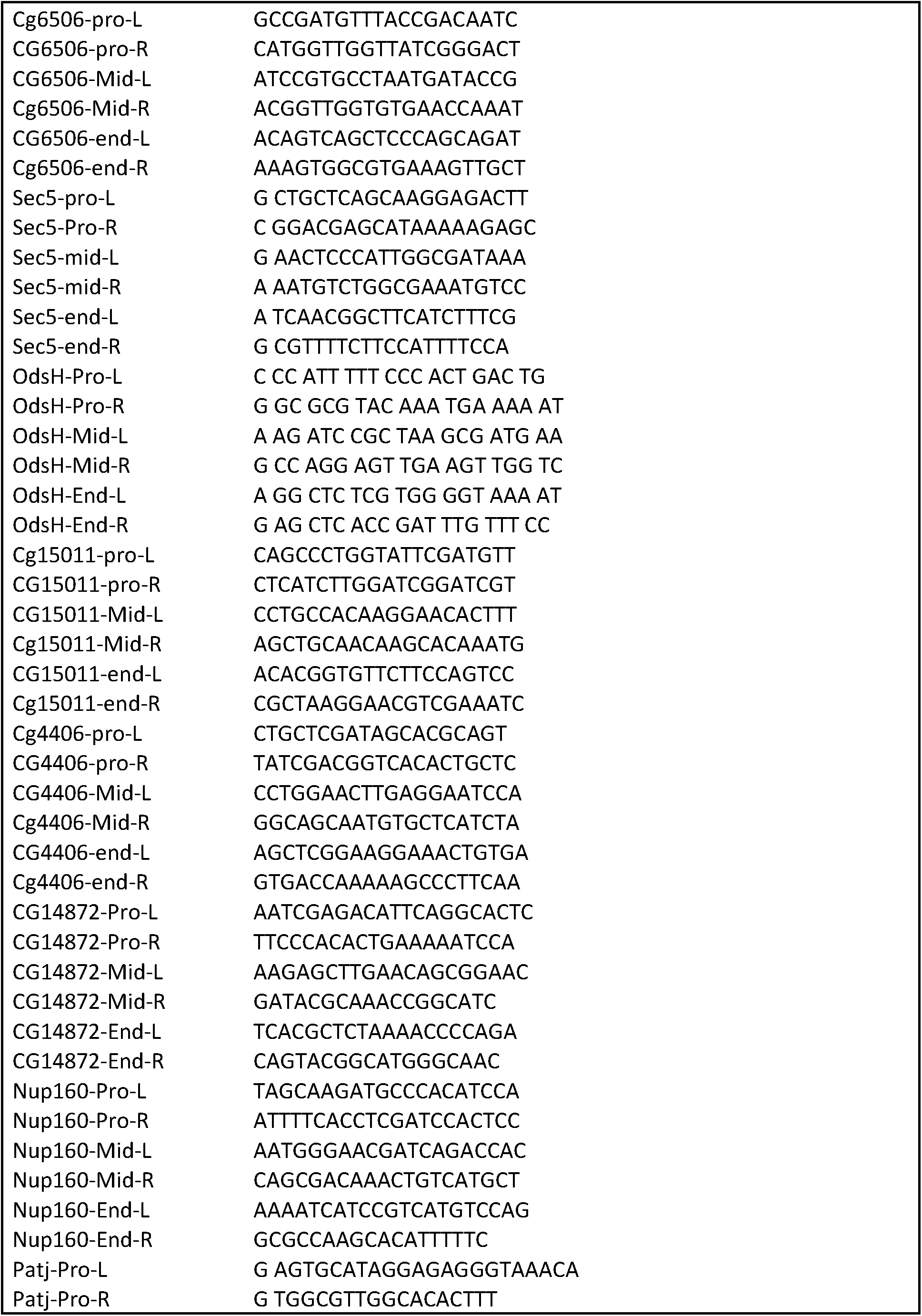
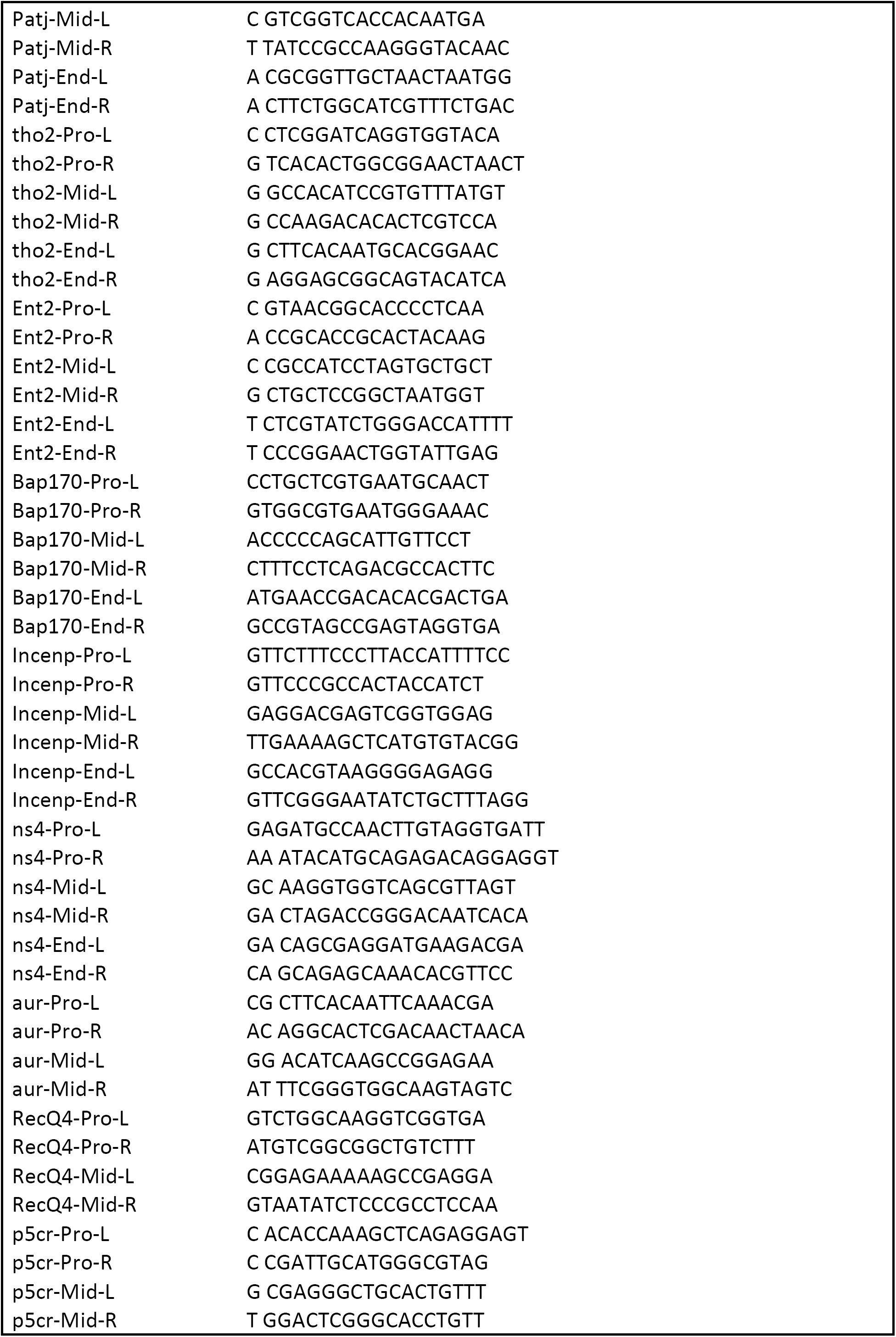
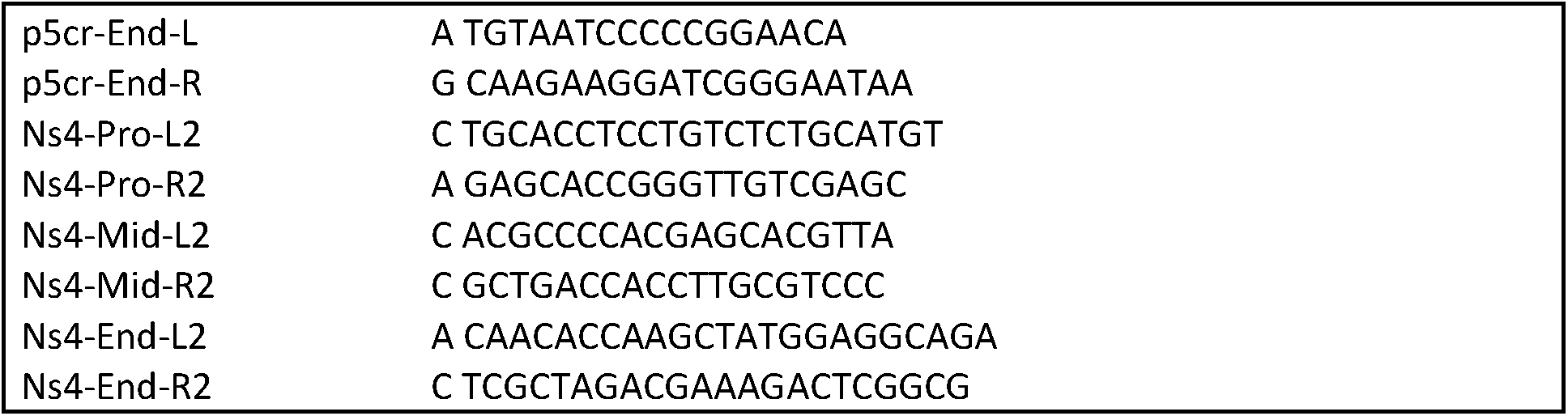
List of primers used for quantitative PCRs

This file contains:

7 Supplementary figures and figure legends

2 Supplementary tables

STAR Methods

Key Resources

### STAR Methods

## Experimental Model and Subject Details

### Drosophila cell cultures

S2 cells and Kc cells were cultured in Schneider’s medium (Gibco) supplemented with 10% FCS and 0.05% Pluronic F-68 (Sigma-Aldrich). The cultures were maintained on shaking incubators at 27 °C, at a speed of 80 rpm. The cells were kept at a density of 1-16 million/ml.

### *Drosophila* rearing conditions and genetics

Unless otherwise specified, flies were reared on a standard cornmeal fly medium at 25 °C, 70% relative humidity and 12 hr dark/12 hr light cycle. The following stocks were obtained from the Bloomington stock centre or kindly donated:

y1 w*; P{tubP-GAL4}LL7/TM3, Sb1 (Bloomington stock #5138)

w1118;P{da-GAL4.w-}3 (Bloomington stock #8641)

y1 w1118; P{lacW}nsl1j2E5/TM3, Sb1 (Bloomington stock #12304) w1118; P{GD13852}v24248 (UAS-nsl3dsRNAi) (VDRC stock #24248)

y1 v1; P{TRiP.JF01299}attP2 (UAS-Nurf-38dsRNAi) (Bloomington stock #31341)

y1 sc* v1; P{TRiP.HMS00065}attP2 (UAS-Nurf-301dsRNAi) (Bloomington stock #33658)

y1 sc* v1; P{TRiP.HMS00628}attP2/TM3, Sb1 (UAS-IswidsRNAi) (Bloomington stock #32845)

y1 v1; P{TRiP.HMC03329}attP40 (UAS-Mi-2dsRNAi) (Bloomington stock #51774)

y1 w1118; nsl1e(nos)1/TM6B, Tb (Yu et al., 2010)

All lines used in this study were generated via standard genetic crosses from the above listed stocks. To assess the genetic interaction between nsl1 and nsl3, Nurf-38 and Nurf-301, w;; daGal4/TM6Tb and w;; daGal4, nsl1j2E5/TM6B, Tb virgin females were crossed with UAS-nsl3dsRNAi, UAS-Nurf-38 and Nurf-301 males and flies were allowed to develop at 25 °C or 29 °C in the latter case. To determine the relative viability, male and female adult flies from at least three independent crosses were counted every other day for a period of 10 days from the start of the eclosion. The total number of non-Tb males and females was divided by the total number of Tb males and females respectively, which were used as an internal control with 100% viability.

To assess the genetic interaction between nsl1 and Iswi and Mi-2, w;; tubGal4/TM6B, Tb, Ubi-GFP and w;; tubGal4, nsl1e(nos)1/TM6B, Tb, Ubi-GFP virgin females were crossed to UAS-IswidsRNAi/TM6B, Tb, Ubi-GFP and UAS-Mi-2dsRNAi males and flies were allowed to lay eggs on yeast-supplemented apple juice plates. 0-6 hour collections were used 24 hrs later to separate fluorescent from non-fluorescent larvae under a fluorescent stereomicroscope and at least 3 plates with 100 larvae/plate were monitored and the rate of pupariation/number of pupae determined over a period of 3 weeks.

It is important to note that although the used heterozygous nsl1 null mutant alleles do not affect the viability of the flies (data not shown), it has been previously reported that nsl1e(nos)1/+ has reduced NSL1 function rendering it amenable to genetic interaction analysis (Yu et al., 2010). When a strong ubiquitous driver, tubGal4, was used for dsRNAi induction at 25°C, this resulted in 100 % lethality (data not shown) recapitulating the loss-of-function phenotype previously reported for nsl3, Nurf-38, Nurf-301, Iswi and Mi-2.

The used combinations of daGal4 or tubGal4 and 29°C, 25°C or 18°C aimed at partial lethality or sensitized genetic background allowing for enhancers or suppressors of the lethality phenotype to be identified.

For Fig. 2D, in order to minimize background effects (Chari and Dworkin, 2013), we chose a set of *UAS-RNAi* lines (Flockhart et al., 2006; Hu et al., 2017; Perkins et al., 2015) that efficiently silence the target genes, leading to 100% lethality, and established loss-of-function mutants of these genes. We used either a weaker Gal4 driver (*da-Gal4*), or lower temperatures, to establish conditions of partial lethality.

## Method Details

### Preparation of nuclear extract from S2 cells

Wild type S2 cells or S2 cells stably expressing the NSL complex members were harvested and washed with ice-cold PBS. The volume of the cell pellet was estimated. The cells were re-suspended and incubated in 5 pellet cell volumes (PCV) of cold hypotonic buffer (10 mM HEPES pH 7.9, 1.5 mM MgCl_2_, 10 mM KCl and Roche complete Protease Inhibitor Cocktail) on ice for 15 minutes. NP-40 was then added to the hypotonic buffer to a final concentration of 1% and the cells were immediately vortexed for 30 seconds. The cells were centrifuged at 2000g for 5 minutes. The supernatant was kept as cytoplasmic extract while the pellet was kept as nuclei. The nuclei were re-suspended and washed with 5 PCV of cold isotonic solution (25 mM HEPES pH 7.6, 2 mM MgCl_2_, 3 mM CaCl_2_, 300 mM sucrose and Roche complete Protease Inhibitor Cocktail). The washed nuclei were carefully re-suspended in 5 PCV of HEMGT 250 buffer (25 mM HEPES pH 7.4, 0.1 mM EDTA, 250 mM NaCl, 1 mM MgCl_2_, 0.1% Triton X- 100, 10% glycerol and Roche complete Protease Inhibitor Cocktail) and rotated for 2 hours at 4 °C. The nuclei were finally centrifuged at 18000g for 30 minutes. The resulting supernatant contained the nuclear extracts.

### Co-immunoprecipitation

The nuclear extract prepared as aforementioned was diluted with an extraction buffer without salt to lower the NaCl concentration from 250 mM down to 150 mM. If the extract went through a freeze-thaw process, it would always be further centrifuged at 14000 g for 15 minutes before being used in immunoprecipitation experiments. 200 ul of the extract was used for immunoprecipitation. HEMGT 150 buffer (25 mM HEPES pH 7.4, 0.1 mM EDTA, 150 mM NaCl, 1 mM MgCl_2_, 0.1% Triton X-100, 10% glycerol and Roche complete Protease Inhibitor Cocktail) was used to increase the volume to 600 ul. Protein A/ protein G beads (Protein A/G Sepharose 4 Fast Flow from GE Healthcare) was used to pre-clear the extract. Before use, all beads were pre-equilibrated with HEMGT 150. Extracts were incubated with 2 to 5 ul of the antiserum (depending on the strength of the antibody) for 4 hours to overnight at 4 °C. 5 ul of rabbit/rat IgG or pre-immune serum was used in the control experiments. 40 ul of protein A/ protein G slurry was added and the tubes were mixed on a rotator at 4 °C for another hour. The beads were then spun down at 1000 g for 1 minute. 6 HEMGT- 150 washes with 1 minute of rotation in 4 °C were performed. Finally, the beads were boiled in 100 ul of 2X Laemmli Sample Buffer (Rotiload, Roth) for 5 minutes. For immunoprecipitated samples, 18 mM of iodoacetamide was added to block reduced cysteine residues and ensure complete denaturation of the eluted antibodies. The samples were incubated for 30 minutes in the dark. 15 ul of the samples was analyzed using SDS PAGE. Anti-ISWI, anti-BRM and anti-Mi-2 (from Dr. Peter Verrijzer), anti-CHD3 antibody (from Dr. Alexander Brehm), anti-NURF38 (from Dr. Paul Badenhorst) are used for western blot analysis. Anti-NSL complex antibodies (Mendjan et al., 2006) and anti-NURF301 are the same antibodies as in the ChIP experiments.

For co-immunoprecipitation experiments using stable cell lines, anti-FLAG M2 agarose beads (Sigma) were used. The beads were mixed with the extract for 2 hours at 4 °C on a rotating wheel.

### Knockdown experiments in S2 cells

The double-stranded RNAs against the NSL complex subunits and chromatin remodelers were designed to complement the exon sequences of the respective proteins with a length of 250 to 350 bp. These double-stranded RNAs did not show complementarity to other genes (with 18 bp seeds) besides the genes of interest which was determined by E-RNAi {Horn, 2010 #50}. Double-stranded RNAs against the GST or GFP sequence were used as a control.

Since the double-stranded RNAs were targeting the exons of the genes, genomic DNA was used as a template in the PCR amplification. For double-stranded RNAs against GST, the plasmid pET-41a (milipore), which contains a GST tag, was used as a template. The amplification was done for 35 cycles using Phusion High Fidelity Polymerase according to the manufacturer’s recommendations. The DNA was loaded on an agarose gel to ensure that a specific PCR product was amplified. When multiple bands appeared on the gel, a different primer pair was used in the amplification, or the DNA was purified by gel extraction. The DNA could also be purified with QIAquick PCR Purification Kit (Qiagen) if the PCR reactions were confirmed to produce one specific product. Double-stranded RNAs were generated using the Ribomax Large Scale T7 *in vitro* transcription system (Promega) according to the manufacturer’s instructions. The RNAs were then purified using the MEGAclear (Ambion) and eluted in nuclease-free water (Invitrogen). To ensure that the two strands of the RNAs annealed perfectly to each other, the RNAs were heated to 65 °C for 10 minutes and allowed to cool down slowly to room temperature.

For knockdown experiments, 2 ml of S2 cells at 1 million cells per ml was plated in 6-well dishes and left to attach for 1 hour. 10 ug of purified double stranded RNA was diluted with Schneider’s medium. The RNA transfection was performed using Lipofectamine RNAiMAX Reagent (Invitrogen), according to manufacturers instructions. The double stranded RNAs were mixed with the transfection reagent and kept at room temperature for 15 minutes before the mixture was added to the cells. The cells were harvested after 4 days and cell numbers were counted.

### Luciferase assays

100 ul of cells at 1 million cells per ml was plated on 96 wells plates. Cells in each well were transfected with a plasmid mixture of (1) 200ng of pG5luc, which contains the firefly luciferase gene whose expression is controlled by UAS sequences (2) 2ng of pRL-hsp70, which contains a constitutively expressed Renilla luciferase gene and (3) 50ng of pAc5.1 vector containing either Gal4DBD-tagged NSL3 protein, or Gal4DBD alone. The Renilla luciferase acts as a control for normalization of transfection and cell number variation. Transfections were carried out with X-tremeGENE DNA Transfection Reagents (Roche). After 2 days of incubation, cells were lysed (Dual-Luciferase Kit, Promega) and luminescence was measured by using Mithras plate reader (Berthold). To test the Gal4-NSL3 activity upon depletion of the NSL complex and chromatin remodelers, double stranded RNAs against the proteins of interest were transfected to the cells on 96 well plates. The knockdown experiments were carried out as described in the previous section. 2 days after the RNA transfection, the luciferase gene-containing plasmids were transfected to the cells as described above.

### Chromatin Immunoprecipitation (ChIP)

50 million S2 cells were harvested and washed with PBS. The cell was resuspended in 1ml of crosslinking solution (50 mM HEPES, 100 mM NaCl, 1 mM EDTA, 0.5 mM EGTA), 37% formaldehyde was added to a final concentration of 1.8%. The tube was rotated at room temperature for 10 minutes and the reaction was quenched with 2.5 M glycine at a final concentration of 125 mM. The cells were spun down immediately after addition of glycine, and the solution was replaced with fresh PBS with 125 mM glycine. The cells were resuspended and rotated for 5 minutes at room temperature. All the steps that follow were carried out either at 4 °C or on ice. The cells were pelleted by centrifugation at 2000rpm for 5 minutes. The cells were washed three times with Paro 1 (10 mM Tris pH 8.0, 10 mM EDTA, 0.5 mM EGTA, 0.25% TritonX-100), four times with Paro 2 (10 mM Tris pH 8.0, 200 mM NaCl, 1 mM EDTA, 0.5 mM EGTA) and twice with RIPA (140 mM NaCl, 25 mM HEPES pH 7.5, 1 mM EDTA, 1% TritonX-100, 0.1% SDS, 0.1% DOC). For each washing step, cells were resuspended in the mentioned buffers and rotated in the cold room for 5 minutes, before they were spun down again. After all the washing steps, cells were resuspended in 500 ul RIPA and sonicated using a Branson sonificator with the following settings: power output: 3, Duty cycle: 40, cycle: 20 seconds on, 50 seconds off. In total, 35 cycles of sonication were performed. After sonication, an aliquot of the chromatin was decrosslinked, treated with Protease K/RNaseA and the DNA was purified. The DNA was analyzed with agarose gel to ensure that the chromatin was sheared into fragments with average length of 200 bp. The number of cycles can be adjusted to achieve the desired fragment length.

For NSL1 and NURF301 MNase-ChIP-seq, we used MNase for chromatin shearing. Cells were washed as above described and suspended in MNase digestion buffer. A small aliquot of the samples is used to determine the optimal MNase concentration (20-200 U/ml tested) to obtain the desired fragment size. Digestion is performed in room temperature for 20 minutes and finally quenched by addition of 1 mM EDTA and 2X RIPA buffer.

Chromatin was centrifuged at 14000 rpm for 30 minutes and the pellet was discarded. The supernatant was incubated with Protein A/ Protein G sepharose beads and rotated in the cold room for 30 minutes to preclear the chromatin. The concentration of the chromatin was determined with a NanoDrop 1000 Spectrophotometer (thermo). For each IP, 10 ug of chromatin was diluted in RIPA to fill up the volume to 500 ul. 3 ul of antibody was added to the chromatin and rotated overnight at 4 °C. Anti-NURF301 (Novus Biologicals) and anti-NSL1 (Mendjan et al., 2006) antibodies are used. Protein A/G sepharose was blocked with 0.1% BSA and 1 mg/ml of short dsDNA (15 bp) for 1 hour. 20 ul of sepharose was added to the chromatin/antibody mixture and incubated for 3 hours at 4 °C. Protein A sepharose was used if the antibodies came from rabbit or guinea pig; protein G agarose was used if the antibodies came from rat. The sepharose bead was then washed six times for 5 minutes in RIPA buffer, once in LiCl buffer (0.25 M LiCl, 10 mM Tris pH 8, 1 mM EDTA, 0.5% NP-40, 0.5% DOC), and two times in TE buffer. The bead was incubated at 65 °C overnight in TE buffer to reverse the crosslink. The sample was then treated with RNaseA (0.2 mg/ml) for 30 minutes at 37 °C, and with Proteinase K (0.05 mg/ml) for 2 hours at 50 °C. Finally, DNA was purified using Minelute columns (Qiagen) and eluted in nuclease-free water. This protocol generated enough material for quantitative real time PCR analysis. For ChIP-seq analysis, the same protocol was scaled up until 10 ng of DNA was achieved.

The eluted DNA from the ChIP protocol was analyzed by qPCR using SYBR Green Master Mix (Roche) and ABI7500 real-time PCR thermocycler (Applied Biosystems, Inc.). For ChIP samples, 10% and 1% INPUT materials were included in the analysis. Efficiency of the ChIP (as input recovery) was determined as the amount of immunoprecipitated DNA relative to input DNA.

### Micrococcal nuclease digestion followed by sequencing (MNase-seq)

50 million S2 cells were harvested and washed with PBS. Cells were counted very carefully to ensure that the same number of cells was collected for each sample. The cells were resuspended in 1 ml of crosslinking solution (50 mM HEPES, 100 mM NaCl, 1 mM EDTA, 0.5 mM EGTA) and 37% formaldehyde was added to a final concentration of 1%. The tube was rotated at room temperature for 10 minutes and the reaction was quenched with 2.5 M glycine at a final concentration of 125 mM. The cells were spun down immediately after addition of glycine and the solution was replaced with fresh PBS that contained 125 mM glycine. The cells were resuspended and rotated at room temperature for 5 minutes. The cells were pelleted into equal aliquots by centrifugation with 2000 rpm for 5 minutes. Aliquots were then snap frozen in liquid nitrogen.

Cells were permeabilized with NP-40 as described in the previous section. The chromatin was digested with various amount of MNase at 25 °C for 10 minutes. Digested chromatin was then analyzed on an agarose gel to reveal the digestion pattern. We aimed to achieve a slightly under-digested condition with di- and tri-nucleosomes still visible. In addition, the digestion pattern of each sample needed to be exactly the same. After the optimal amount of MNase was determined for each individual experiment, a new aliquot of the cells was again digested with MNase at 25°C for 10 minutes. The reaction was stopped by addition of EDTA, NaCl and SDS to an end concentration of 20 mM, 150 mM and 1% respectively. The samples were placed on ice during handling to ensure that the MNase digestion was halted completely. The chromatin was then de-crosslinked overnight at 65 °C overnight in TE buffer. The sample was treated with RNaseA (0.2 mg/ml) for 30 minutes at 37 °C, and with Proteinase K (0.05 mg/ml) for 2 hours at 50 °C. Finally, DNA was purified using Minelute columns (Qiagen) and eluted in nuclease-free water.

The eluted DNA was then analyzed with the E-Gel Pre-cast Agarose Gels system (Invitrogen). Bands at positions corresponding to 150 bp, 300 bp, 450 bp, represented mono- di and tri-nucleosomes, respectively. The 150 bp fragment was purified from the E-gel according to manufacturers’ specifications and used to generate the library for the Illumina paired-end sequencing. To minimize the effect of PCR artefacts, only 3 cycles of PCR were performed in the library generation procedure.

### MAPCap

Protein G magnetic Dynabeads (Invitrogen) were prepared with IPP buffer (50 mM Tris-HCl pH 7.4, 150 mM NaCl, 0.1% NP-40). 2.5 µg of anti-m7G antibody (SYSY, catalog nr: 201 011) were incubated with the beads for at least 1 hour in 4°C. The beads were finally washed twice with IPP buffer. RNA extractions were carried out with the Direct-zol Miniprep kit (Zymo Research). Abundant small RNAs (< 200 nt) were removed using the RNA Clean and Concentrator Kit (Zymo Research) and eluted in 100 µl TE buffer (10 mM Tris-HCl pH 8.0, 1 mM EDTA). RNA fragmentation was performed using the Covaris E220 Focused-ultrasonicator (Duty cycle: 10%, Intensity: 5, Power: 175 W, Cycles/Burst: 200, time: 140 s). After sonication, the capped RNA was captured with the antibody-coupled Protein G magnetic beads for 1 to 2 hours in 4 °C. Then, the beads were washed 3 times with IPP buffer. RNA 3’ ends were repaired using PNK. Custom-made, barcoded adapters were ligated to the RNA using T4 RNA Ligase 1 for 1 h at 25 °C. Excess adapters were washed away with IPP buffer and RNA was purified by column purification. Isolated RNA was reverse-transcribed and treated with RNase H. cDNA was column-purified and circularized with CircLigase for 2–16 hours. Circularized cDNA was directly PCR amplified, the amplified library was finally cleaned up using Ampure beads.

## Quantification and Statistical Analyses

### MNase-seq analysis

#### MNase-seq read mapping

The paired-end reads were aligned using Bowtie2 (version 2.0.5; (Langmead and Salzberg, 2012)) or with STAR (version 2.5.1b; (Dobin et al., 2013)) against the *Drosophila melanogaster* reference genome (version dm3, UCSC). Default parameters were used, except for the STAR alignment, where we set the option alignIntronMax to 1. In case of Bowtie2 resulting BAM files were sorted using samtools (Li et al., 2009), while this step was omitted for STAR alignments as they were already sorted. The sorted BAM files were indexed using samtools.

#### Estimation of the nucleosome signal

To get an unbiased estimate of the nucleosome signal we counted the 5’ starting positions of the reads separated by their orientation, i.e. plus- and minus-strand alignments. This results in two vectors **p** and **m** with components *p*_*i*_ and *m*_*i*_, where *i* indicates the position and **p** corresponds to the counts for plus-strand reads and **m** for minus-strand reads. For all positions *i* where *p*_*i*_ *+ m*_*i*_ > 0 we calculated *x*_*i*_ = (*p*_*i*_ - *m*_*i*_)/(*p*_*i*_ *+ m*_*i*_), while *x*_*i*_ = 0 for positions *i* where *p*_*i*_ *+ m*_*i*_ = 0.

For a well-positioned nucleosome at its dyad position *k* we expect many more plus-strand reads than minus-strand reads at its left border at position *k* - 73 and vice versa many more minus strand reads than plus-strand reads at its right border at position *k +* 73. To account for the inprecise mapping of the nucleosome borders by MNase-seq, we defined a function 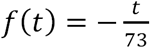 for −73 ≤ *t* ≤ 73 and zero otherwise. This function was convolved against the vector **x** to obtain the nucleosome signal 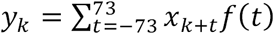 at the nucleosome dyad position *k*. This nucleosome signal is between −74 and +74, where a negative value indicates depletion and positive signal enrichment. Note, that this procedure yields a normalized nucleosome signal, which is independent on library size.

### ChIP-seq analysis

#### ChIP-seq read mapping

The ChIP-seq reads were aligned either with Bowtie2 or STAR and processed as outlined in the MNase-seq read mapping section.

#### Peak calling

Peaks were called using the Bioconductor package normR (https://www.bioconductor.org/packages/normr; (Kinkley et al., 2016). Reads/fragments were counted using the Bioconductor package bamsignals (https://www.bioconductor.org/packages/bamsignals).

For single-end ChIP libraries the fragment size was estimated by convolving the vector **m** of counts of minus strand reads against the corresponding vector **p** of plus strand reads. The fragment size was determined by choosing the distance *k* for which *Σ p*_*i*_*m*_*i+k*_ was maximal.

For paired-end ChIP and input libraries we counted the number of fragment midpoints falling into non-overlapping 200 base pair bins along the genome. For single-end ChIP and input libraries the 5’ positions of the reads were shifted by half the fragment size to the right for plus strand and to the left for minus strand reads. For each ChIP experiment a normR analysis was performed using the function enrichR, where the count vectors for the ChIP and the corresponding input control served as input.

Normalized log_2_ Chip over control were calculated by *f* = log_2_(y *+* a) - log_2_(*x +* b) *+* log_2_(b) - log_2_(a), where y corresponds to the number of ChIP fragments/reads in a bin of size *w*, x to the number of control fragments/reads in the same bin. a to the average number of ChIP fragments/reads in 200 base pair bins that had a qvalue > 0.1 divided by 200 and multiplied by *w*, and b to the average number of control fragments/reads in 200 base pair bins that had a qvalue > 0.1 divided by 200 and multiplied by *w*.

### RNA-seq analysis

#### RNA-seq read mapping

The paired-end RNA-seq reads were aligned using STAR (version 2.5.1b) against the *Drosophila melanogaster* reference genome (version dm3, UCSC) plus the *Saccharomyces cerevisiae* genome (release R64-1-1, ENSEMBL) including the corresponding gene annotations from both species.

#### DEseq2 analysis

Standard RNA-seq analyses are not able to detect global shifts in gene expression. We expected that a NSL1 knockdown may lead to a global downregulation of almost every expressed gene. In such a scenario, genes that are un- or only mildly affect by the NSL1 knockdown would appear upregulated. To mitigate this problem, we spiked yeast RNA into the RNA preparations according to the cell number, which had been estimated by qPCR. The yeast RNA serves as external control to normalize the RNA-seq data, because the abundance of yeast RNAs remains unchanged. Thus, after mapping to the combined *Drosophila* and yeast reference genomes, we obtained the read counts for the 16 yeast chromosomes. And used these counts to calculate size factors using the DESeq2 (Love et al., 2014) function estimateSizeFactorsForMatrix using triplicates for control, MOF, NSL1 and NURF knockdowns.

For each condition and replicate we counted reads mapping to annotated exon ranges per gene, where overlapping exons were merged. We counted only the first mate to retain the strand information.

A preliminary principal component analysis on the rlog transformed counts revealed that two samples, one control and one NSL1 sample, were outliers, i.e. did not cluster with their respective conditions. These were removed prior to any downstream analysis.

With the remaining 10 samples we performed a DESeq2 analysis, where we manually set the size factors to the ones estimated by the yeast spike ins. We used the condition, i.e. either control, MOF, NSL1 or NURF knockdown, as explanatory variables. After the DESeq analysis we obtained results by contrasting MOF, NSL1 or NURF against the control.

### gDNA Analysis

#### Genomic DNA immunoprecipitation followed by sequencing

Genomic DIP–seq experiments were performed as described (Gossett and Lieb, 2008) with few modifications. The pellet from 1 × 10^8^ S2 cells was suspended in 1 ml of lysis buffer (10 mM Tris pH 8, 25 mM EDTA, 100 mM NaCl, 0.1% SDS) and treated with protease K overnight at 55 °C. The DNA is tested with Bradford assay and protein gel to ensure that no proteins remain. DNA is sonicated with Covaris sonicator (power 175 W, duty factor 10%, cycles per burst 200, 250 s) to obtain fragment size average of 200-bp. DNA was purified with the DNA Clean & Concentrator (Zymo).

FLAG-NSL3 and FLAG-GFP were expressed in baculovirus. The extract was prepared two days after infection. Details of the protocol are available upon request. Proteins were purified using FLAG bead slurry (sigma). The beads were incubated with the extracts for 2 hours followed by 4 washes in HEMGT buffer (25 mM HEPES, pH 7.9, 12.5 mM MgCl2, 150 mM KCl, 10% glycerol). DIP-seq experiment was performed with 1µg of genomic DNA and 1.5 µg of FLAG-NSL3 or FLAG-GFP at 4 °C for 2 hours in 500 µl of DIP buffer (2 mM Tris-HCl pH 7.5, 100 mM KCl, 2 mM MgCl2, 10% glycerol, 10 µM ZnCl2). The FLAG bead was washed six times with 1ml of DIP buffer. Elution was performed with 3X FLAG peptide (100µg/ml, overnight) or digestion with proteinase K (1 h at 56 °C), DNA was purified with the DNA Clean & Concentrator (Zymo). The sequencing was performed using the Illumina HiSeq platform with a 75-bp paired-end kit.

#### gDNA read mapping and peak calling

The paired-end gDNA-seq reads were aligned using STAR (version 2.5.1b) against the *Drosophila melanogaster* reference genome (version dm3, UCSC) and processed as outlined in the MNase-seq read mapping section.

Peak calling of the gDNA pulled down by GFP, MCRS2, NSL1 and NSL3 was performed as outlined in the ChIP-seq peak calling section using 50 base pair bins. As control served the input to the pull down experiments. After the identification of enriched bins, we merged directly adjacent bins and determined the maximal bin within the so-derived regions. The coordinates of these maximal bins per enriched region were referred to as peak center.

### Correlation of gDNA enrichment to sequence features

We computed the normalized and regularized log enrichment using the normR function getEnrichment with parameter standardize = FALSE (Kinkley et al., 2016). Using the peak centers as anchor points, we determined the AT content in 50 base pair bins in a window +/- 1,000 base pairs around these anchor points. We sorted the peaks by the average log enrichment +/- one bin around the peak center bin.

For the whole genome, we repeated the normR analysis using 200 base pair bins. For each of the 200 base pair bins, we calculated the AT content as well as the frequency of all 1,024 possible 5mers. We plotted the AT content in percent against the log enrichments for all bins which were not filtered out by the T-filter employed in normR and calculated the Pearson correlation coefficient.

To exclude the possibility that not the AT content but rather a certain AT-rich motif explains the *in vitro* binding of NSL3 we linearly regressed the *in vitro* NSL3 log enrichment over control against the AT content + the frequency of all possible 5mers. This effectly removes transitive, i.e. indirect correlations. For example, if the frequency of the 5mer AAAAA is correlated to the NSL3 log enrichment only because it is more likely to find such a 5mer in an AT-rich sequence then the partial correlation framework would reduce the information contained in the frequency AAAAA about the NSL3 log enrichment. However, if the frequency of AAAAA remains informative for the NSL3 log enrichment after accounting for its increased frequency in AT-rich sequences then this would be a statistical argument to propose that the AAAAA is preferentially bound by NSL3.

### Prediction of in vivo NSL3 MapCap TSS targets

We used the MapCap TSSs as anchor points to compute the NSL3 enrichment in 61 (30 upstream, 1 corresponding to the TSS, and 30 downstream) 29 base pair bins and the AT content in the same bins. We clustered the MapCap TSSs by the average NSL3 enrichment 11 bins around the TSS via kmeans into NSL3 bound and non-bound. Next, we assigned the value +1 for NSL3 bound TSSs and −1 for NSL3 non-bound TSS. The resulting vector was used as response variable for logistic regression. Because ∼70% of the TSSs were NSL3 bound, we used a sampling strategy, were we sampled the same number of NSL3 bound and non-bound TSSs with replacement to get a balanced set. We then performed standard linear regression using 10-fold cross validation.

We performed a similar analysis with the DRE motif, where we used a window of −70 to −10 (i.e. 61 features) upstream of the TSS. This region should encompass the preferential location of the DRE motif.

To assess the goodness of fit, we randomly sampled class assignments, i.e. NSL3 bound and non-bound, with probability 0.687 for NSL3 bound and 0.313 for NSL3 non-bound 1,000,000 times. For each sample we calculated the number of correctly “predicted” class labels.

We also checked for the occurrence of the core promoter motifs: DRE, TATA, INR, MTE and DPE around the MapCap TSSs. We computed the PWM scores around and assigned a positive match to a binding site by requiring that the score is among the 1% highest scores along the genome.

### MapCap analysis

#### MapCap read mapping

MapCap paired-end reads were mapped as outlined in the RNA-seq read mapping section. Prior to mapping the corresponding unique molecular identifier (UMI) was added to the read name to preserve it for downstream analysis.

#### MapCap analysis

The MapCap approach aims at identifying 5’ cap positions at single base pair resolution. Due to the limited input material, an amplification of the library via PCR is necessary. The PCR leads to the problem that it remains unclear whether an accumulation of reads at a certain 5’ cap position is due to high expression or due to amplification. Furthermore, the PCR step introduces additional variance in the data, by differential amplification rates due to sequence composition and fragment length. To mitigate these problems a UMI was added to each fragment, such that PCR artifacts can be filtered out.

To use the UMI information the UMIs were extracted from the read name and appended as RX tag to the tag section of the BAM record by a custom perl script. PCR duplicates were removed using the Picard (version 2.5.0; http://broadinstitute.github.io/picard/) function MarkDuplicates by setting the options REMOVE_DUPLICATES=true and BARCODE_TAG=RX.

The paired-end MapCap reads can be used to assign a putative 5’ cap position to the corresponding gene. This can be achieved by mapping the second mate to the annotated exons and assigning the 5’ cap position to the corresponding gene. We used a custom perl script to perform these assignments whose output denotes the gene, chromosome, position, strand and count of 5’ caps, which we will refer to as transcriptional start sites (TSSs).

We constructed a union of all TSSs by requiring at least 2 reads per TSS and sample (control, NSL1 knockdowns in triplicates) using a custom perl script whose output denotes the gene, chromosome, position, strand and counts for all samples. To each gene we assigned one TSS, which had the maximal number of summed counts in the three control knockdown samples. From the identity of these TSSs we derived a count matrix, which served as input to a DESeq2 analysis as outlined in the RNA-seq/DESeq2 analysis section.

To identify genes which change their TSS usage, we performed the following analysis: First, we extracted the counts for each TSS of a gene for the control and NSL1 knockdown triplicate samples. Second, we retained only TSSs that were covered either in all three GST or NSL1 KD samples or in all six samples. Third, if more than one TSS remained we calculated the likelihood of two probabilistic models: the Null model, where we used a single multinomial distribution with parameters **θ**^0^ to model the TSSs counts per sample. We parametrized these “success” probabilities by 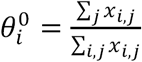, where *x*_*i*,*j*_ denotes the counts for TSS *i* in sample *j*. The alternative model, where we used one multinomial distribution for the control samples and another one for the NSL1 knockdown samples with parameters **θ**^1,control^ and **θ**^1^,NSL^1^, which were parameterized by 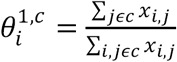, where *c* denotes the condition, control or NSL1.

The likelihood for the Null model for sample *j* is 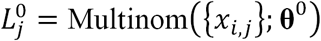 and the likelihood for the alternative model for sample *j* is 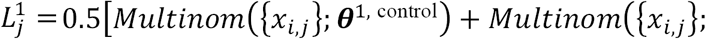 From these per sample likelihoods we computed the total log likelihood by 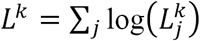 for model *k*.

Finally, we performed a log-likelihood ratio test with |**θ**^0^| - 1 degrees of freedom. We adjust the so-computed p-values for multiple testing by the Benjamini and Hochberg correction.

#### Riboseq analysis

The riboseq data is downloaded from the Gene Expression Omnibus (Dunn et al., 2013). We downloaded the unmapped ribo-seq data, aligned them using Bowtie2 against the *Drosophila* melanogaster BDGP Release 5 (dm3) genome assembly and produced strand-specific coverage to be visualized using Integrative Genomics Viewer (IGV).

### scRNA-seq Analysis

#### Sample preparation

dsRNA-mediated RNAi is performed as described above. Single cells in the control and NSL1 KD samples were carefully diluted, re-suspended and filtered to obtain single cell suspension. The cells are then captured using the 10x genomics Chromium Single Cell 3’ Library and Gel Bead Kit v2, 4 rxns (10x Genomics, 120267) according to the manufacturer’s instructions (Single Cell 3’ Reagent Kits v2 User Guide, CG00052). Sample was loaded to a different lane on a chip (Chromium Single Cell A Chip Kit, 16 rxns, 1000009). After generation of nanoliter-scale Gel bead-in-EMulsions (GEMs), RNA contained in the GEMs were reverse transcribed in a Thermal Cycler (Bio Rad) (53°C for 45 min, 85°C for 5 min, hold at 4°C). Then, the single-strand cDNA was isolated and amplified. Finally, the amplified cDNA was fragmented, end-repaired, A-tailed and indexed using the Chromium i7 Multiplex Kit 96 rxns (120262). The clean-up steps during the library generation was performed with Ampure beads. These libraries were sequenced using the Illumina HiSeq platform with a 75-bp paired-end kit. Alignment and initial processing of sequencing data were performed with CellRanger package using the default settings. The package was used to demultiplex the data, generate FASTQ files, align the sequencing reads to the dm3 reference genome, filter cell and UMI (unique molecular identifier) barcodes.

#### Filtering and Normalization

We used the cellranger Rkit to obtain matrices with cells in the columns and genes in the rows. For each condition (control and NSL1 knockdown) we retained only cells that had more than 2,000 nonzero genes. Next, we selected genes that had nonzero entries in more than 50% of the remaining cells. The filtering resulted in 3,258 cells for the control and 588 for the NSL1 knockdown and in total 1,787 genes.

We implemented a normalization technique, where we iteratively divide the matrix by the outer product between square root of the row and column means. This procedure converges quickly and results in a matrix, whose row means (= genes) and column means (= cells) equal unity. Here, we used 40 iterations.

For the PCA analysis, we concatenated the columns of the control and NSL1 knockdown and performed aforementioned normalization. The resulting balanced matrix was used as input to PCA and we plotted the rotation vectors for the first two principal components, showing that the second principal component is separating control and NSL1 knockdown cells.

To compare variation between conditions, we separately input the control and the NSL1 data. In both cases we obtained a balanced matrix, which we used to calculate the standard deviation for each gene. Note that the mean for each gene is fixed at one, such that the standard deviation for each row of the matrix is equal to the coefficient of variation, which is the standard deviation divided by the mean.

A confounding factor in the estimation of the coefficient of variation is the number of zero and nonzero elements per cell. To control for this factor, we subsampled the control cells by requiring that the distribution of non-zero genes is similar to the one found in NSL1 knockdown. We repeated subsampling 100 times and took the average gene coefficient of variation over these 100 subsamples.

scRNA-seq data was normalized using BASiCS (Vallejos et al., 2015). As input served unique molecular identifier counts for each gene and cell in the control and NSL1 knockdown experiment. To control for technical variability ERCC spike ins were used. We retained only cells that had more than 2,000 nonzero genes and had more than 10 ERCC spike in reads. Thereafter, we selected genes whose mean count was larger than 1 and ERCC spike ins that had at least one nonzero entry. Because there were many more control cells than NSL1 KD cells, we subsampled the control to the same number of cells as in the NSL1 knockdown (560). We repeated this 32 times. After running the BASiCS algorithm, we detected differential variability using the BASiCS_TestDE function. The assignment as a gene that changed variability required that all 32 subsamples flagged the gene as having an increased variability upon NSL1 KD.

### Data and Software Availability

The following datasets have been deposited in the Gene Expression Omnibus repository with the accession number GSE118726:

- MAPCAP – GST RNAi and NSL1 RNAi
- MNase-ChIP-seq with NSL1 and NURF301
- MNase seq – GST RNAi, NSL1 RNAi, INO80 RNAi and NURF301 RNAi
- Genomic DNA-IP-seq with GFP, NSL1, NSL3 and MCRS2
- Single cell RNA-seq – GST RNAi and NSL1 RNAi

### Key Resources

**KEY RESOURCES TABLE**

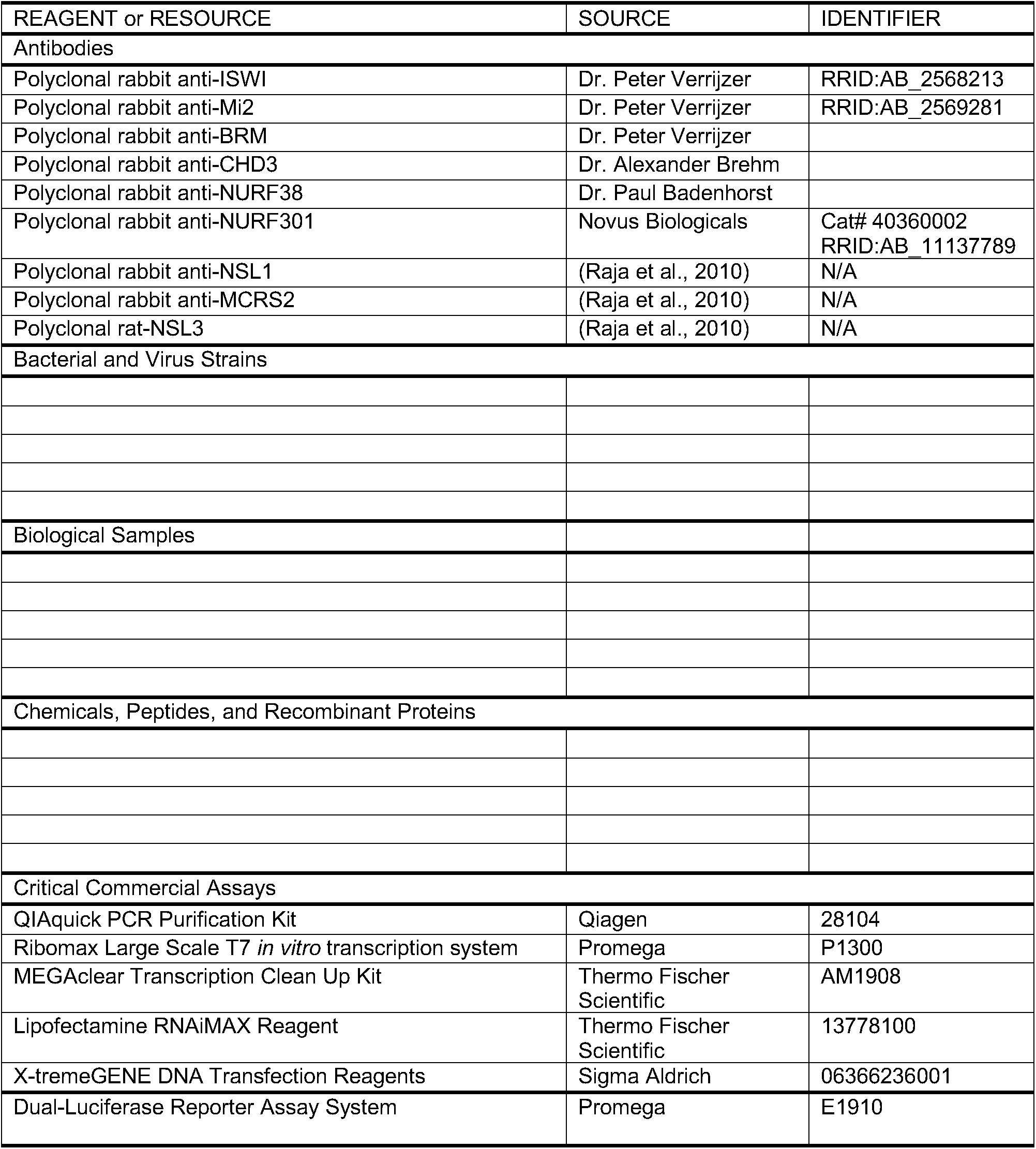

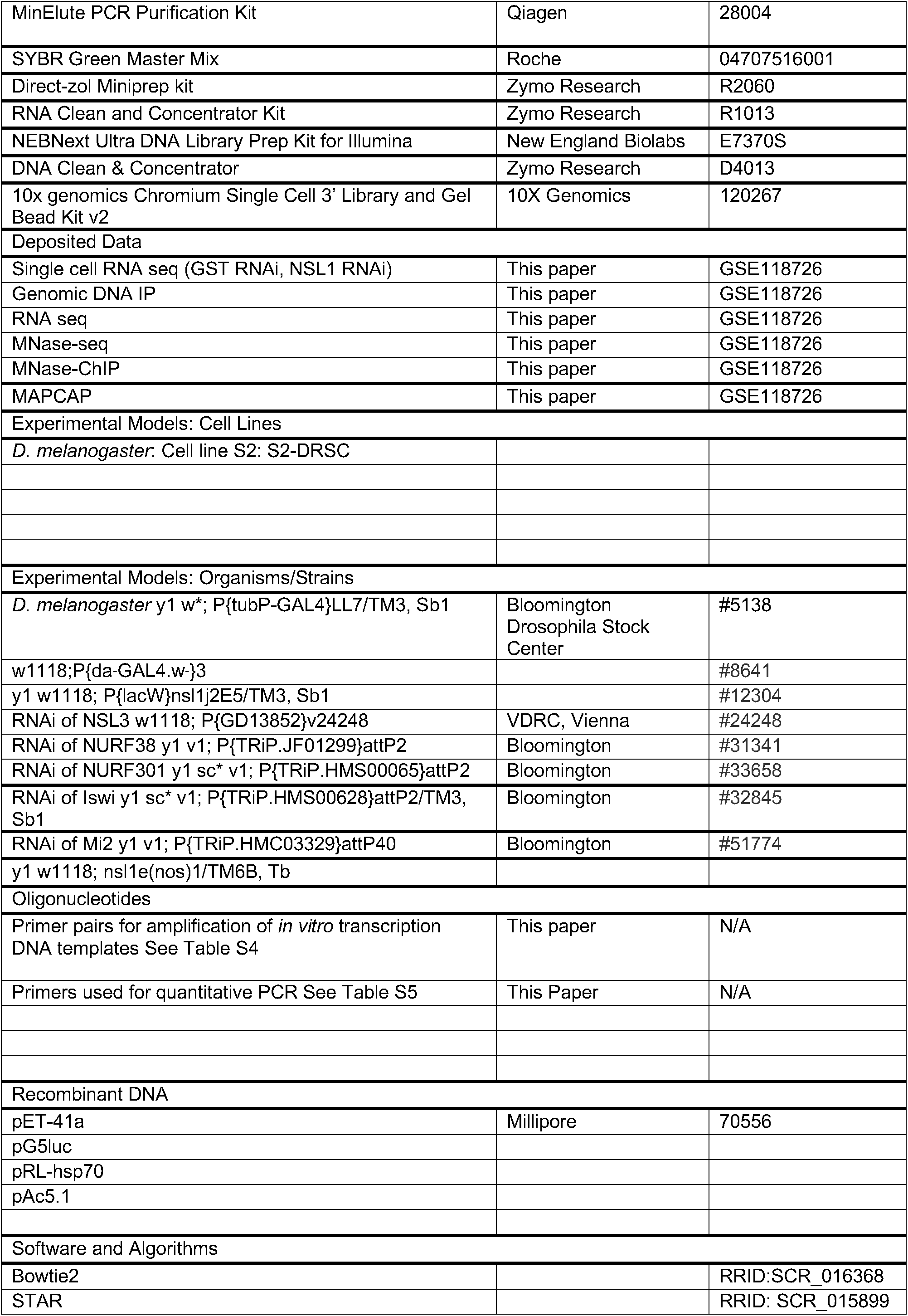

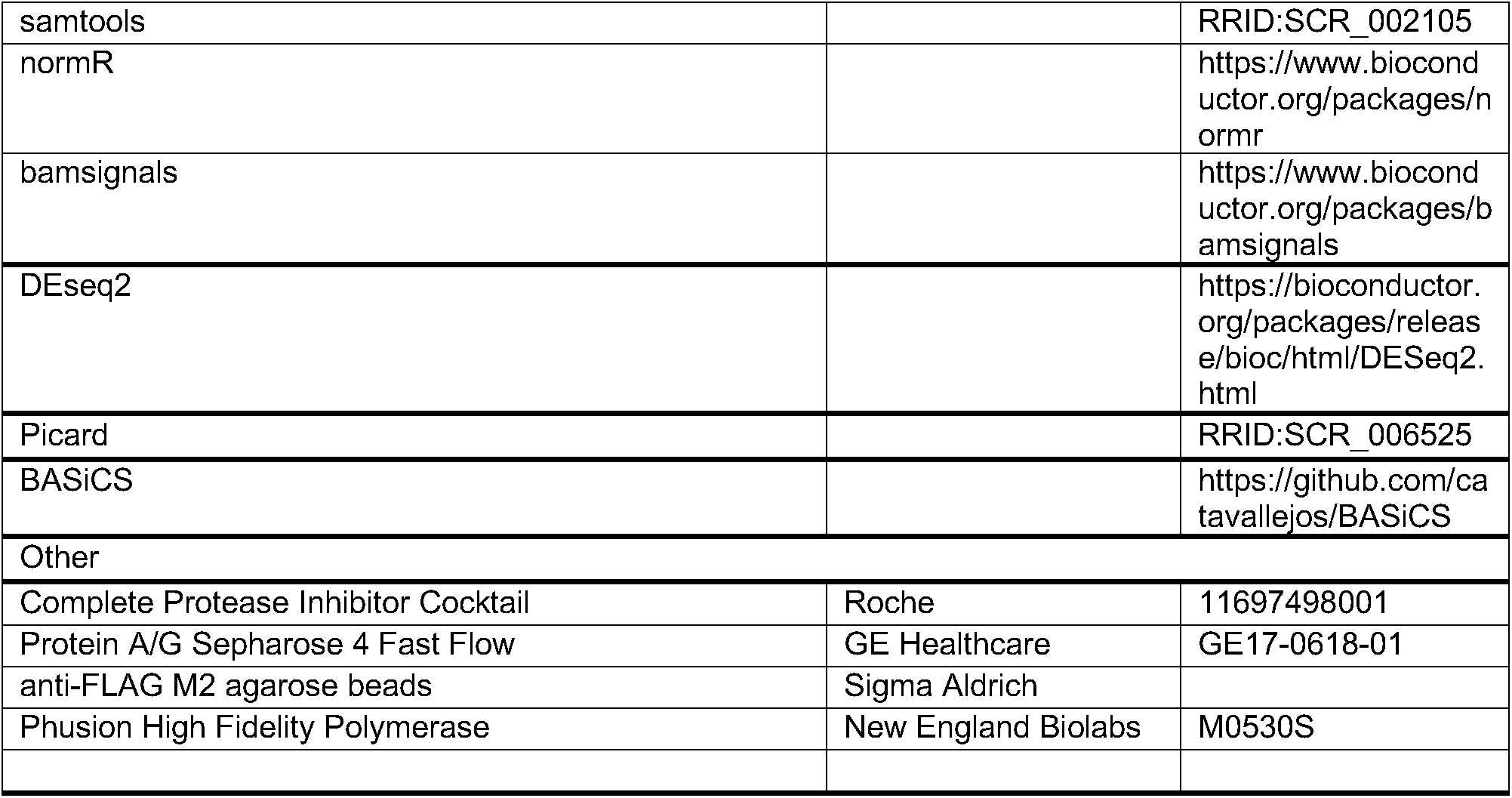

